# Local axonal translation sustains synapse-type-specific neurotransmission

**DOI:** 10.1101/2022.06.27.497827

**Authors:** Hovy Ho-Wai Wong, Alanna J. Watt, P. Jesper Sjöström

## Abstract

Dendritic protein synthesis is a well-known regulator of neurotransmission in the central nervous system. For example, local on-demand protein synthesis in dendrites is required for hippocampal long-term^1,2^ and homeostatic plasticity^3^. Although axonal protein synthesis is a key regulator in neural circuit assembly during early development^4–6^, it is unclear if local translation in mature axons plays a functional role in neurotransmission of the mammalian central nervous system. Here we show that local translation in axons regulates pyramidal cell neurotransmission in primary visual cortex microcircuits in a synapse-type-specific manner^7,8^. Using paired recordings to manipulate pre- and postsynaptic neurons independently, we found that presynaptic translation sustains neurotransmission and regulates short-term plasticity. This was mediated via mTOR and cap-dependent translation, with presynaptic NMDA receptors acting as an up-stream trigger. By isolating the axon from the soma with laser microsurgery, we unveiled that axonal translation was required to sustain neurotransmission. With live endogenous RNA imaging, we found that RNA granules stably docked in individual presynaptic compartments, suggesting bouton-specific regulation. In agreement, presynaptic translation influenced pyramidal cell synapses to neighbouring pyramidal cells, but not to inhibitory Martinotti cells. Our study establishes local protein synthesis in mature axons as a fundamental regulatory principle that governs information transfer at central synapses.

---

Tens of thousands of new protein copies are needed at synapses every minute*^9,10^*. Yet synapses are typically located hundreds of microns from the cell body where, in the textbook view, proteins are made, which raises the question of how proteins are delivered to synapses in a timely manner. Local translation is an attractive solution, as it allows proteins to be synthesised on-site and on-demand^4,6^. Evidence for protein synthesis in dendrites is extensive, ranging from the presence of polyribosomes^4–6,11^ to live visualization of translation^12–16^. Pioneering studies have also demonstrated that dendritic translation regulates neurotransmission^1–3^.

Recent studies^4–6,17,18^ also suggest the existence of local translation in mature axons, i.e., after synapse formation. For example, RNA docking and local protein synthesis occur *in vivo* at presynaptic hotspots^19^, where they may continue to play pivotal roles after circuit assembly^19–23^. Even though hundreds of different mRNAs^24^ and translation machineries^17,24–27^ have been found in mature presynaptic terminals, direct evidence that local translation in mature axons regulates neurotransmission is still lacking. Here, we explore this long-standing question by examining neurotransmission at mature axons from visual cortex layer-5 pyramidal cells (PCs).

To verify that we could acutely inhibit translation in visual cortex slices with cycloheximide (CHX), we assayed translation after 10 minutes of CHX incubation and found that it reliably reduced puromycin incorporation by ~40% (fig. S1a-d). We next explored the effect of CHX on evoked glutamatergic release at layer-5 PC→PC synapses. To efficiently find PC→PC connections, which are sparse^28^, we relied on quadruple patch clamp^29^. We found that within minutes, CHX-mediated inhibition of translation decreased evoked excitatory postsynaptic potentials (EPSPs) in PC→PC pairs (fig. S1e-h).

CHX could reduce EPSP amplitude by blocking protein synthesis presynaptically, postsynaptically, or in a third cell, such as glia (fig. S1k). To examine this, we took advantage of cell-specific pharmacological manipulation in paired recordings. Via the patch pipettes, we loaded two out of four cells in each quadruple recording with the cell-impermeant translation initiation inhibitor m^7^G cap analogue (M7)^1,25,30^ (Fig. 1a). We thus randomly sampled PC→PC connections for which only the pre- or the postsynaptic cell was loaded with M7 (Fig. 1b, c), and found that pre- but not postsynaptic protein synthesis blockade reduced EPSP amplitude (Fig. 1d-h). This demonstrated that presynaptic translation was necessary to sustain synaptic release.

**Fig. 1.**
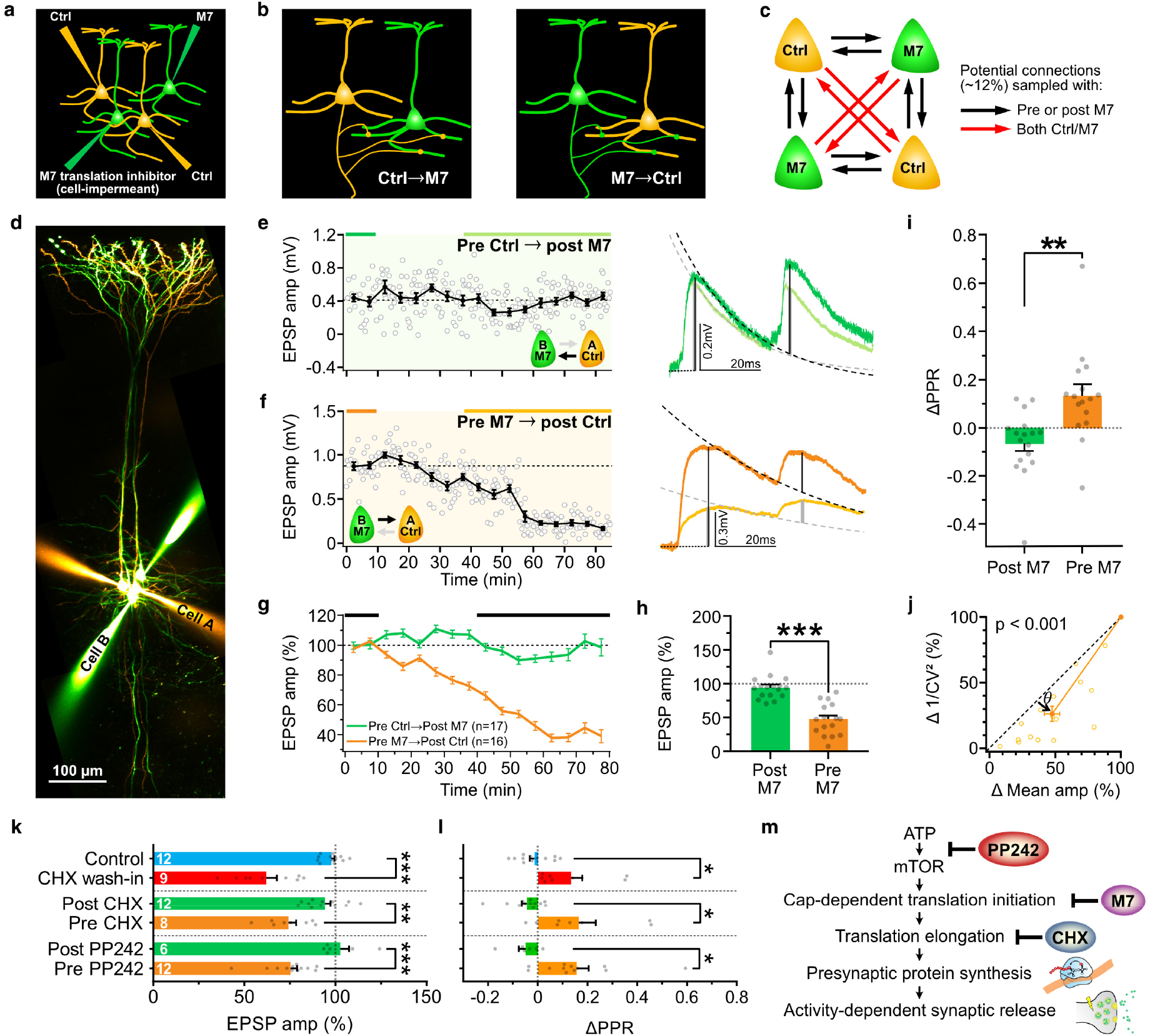
Presynaptic translation sustains activity-dependent neurotransmitter release. **a,** Two out of four cells were loaded with the cell-impermeant translation initiation inhibitor, m^7^G RNA Cap structure analogue (M7), via patching pipettes. **b,** PC→PC connections with only presynaptic M7 or postsynaptic M7. **c,** Quadruple whole-cell recordings were used to compensate for the low connectivity of neighbouring layer-5 PCs^28^. **d,** Sample quadruple recording with reciprocal connections between control cell A and M7-loaded cell B. **e,** For postsynaptic M7, EPSPs were stable. Traces are averaged from and color-coded by the baseline and “after” periods as indicated by bars above the line graph. **f,** With presynaptic M7, EPSPs were suppressed. **g,** Ensemble averages show robustness of finding. (*n = connections*). **h,** EPSP amplitude was suppressed by pre- but not postsynaptic M7 loading (*t_31_ = 6.50*). **i,** Pre- but not postsynaptic M7 increased paired-pulse ratio (ΔPPR) (*t_31_ = 3.39*) in keeping with reduced release. **j,** For coefficient of variation (CV) analysis, points below diagonal for presynaptic M7 (*Wilcoxon test, θ = 10° ± 2°, P < 0.001*) indicates that EPSP suppression was associated with lowered release^31^. **k,** EPSPs after CHX wash-in (*t_9.6_ = 5.99*), CHX loading (*t_18_ = 3.84*) or PP242 loading (*t_16_ = 4.33*). **l,** ΔPPR after CHX wash-in (*t_10.9_ = 2.99*), CHX loading (*t_8.3_ = 2.99*) or PP242 loading (*t_16_ = 2.87*). **m,** Schematic of drug actions on translation pathway and on release. Mean ± s.e.m. \**P* < 0.05, \*\**P* < 0.01, \*\*\**P* < 0.001. Student’s t test or unequal variance t test.

To further explore how M7 affected neurotransmission, we analysed synaptic response paired-pulse ratio (PPR) and coefficient of variation (CV)^31^ (Fig. 1i, j). We found that pre- but not postsynaptic M7 increased PPR and reduced 1/CV^2^, indicating reduced release. Both methods thus independently revealed that presynaptic protein synthesis sustains PC→PC release. We obtained indistinguishable results by loading CHX or the ATP-competitive mTOR inhibitor PP242 (Fig. 1k, l and figs. S2, S3). In summary, evoked release at PC→PC synapses relies on translation in the presynaptic cell via an ATP-mTOR and cap-dependent translation initiation pathway (Fig. 1m).

To directly visualize synaptic release, we stained vesicles by FM 1-43 dye loading^32^ and live-imaged presynaptic boutons using 2-photon laser scanning microscopy (2PLSM) (Fig. 2a, b). When action potentials (APs) were elicited, PC boutons de-stained due to vesicle release, but not in PCs loaded with the M7 translation inhibitor (Fig. 2c and fig. S4a). Together, our electrophysiology and live-imaging data show that presynaptic translation sustains evoked synaptic release.

**Fig. 2.**
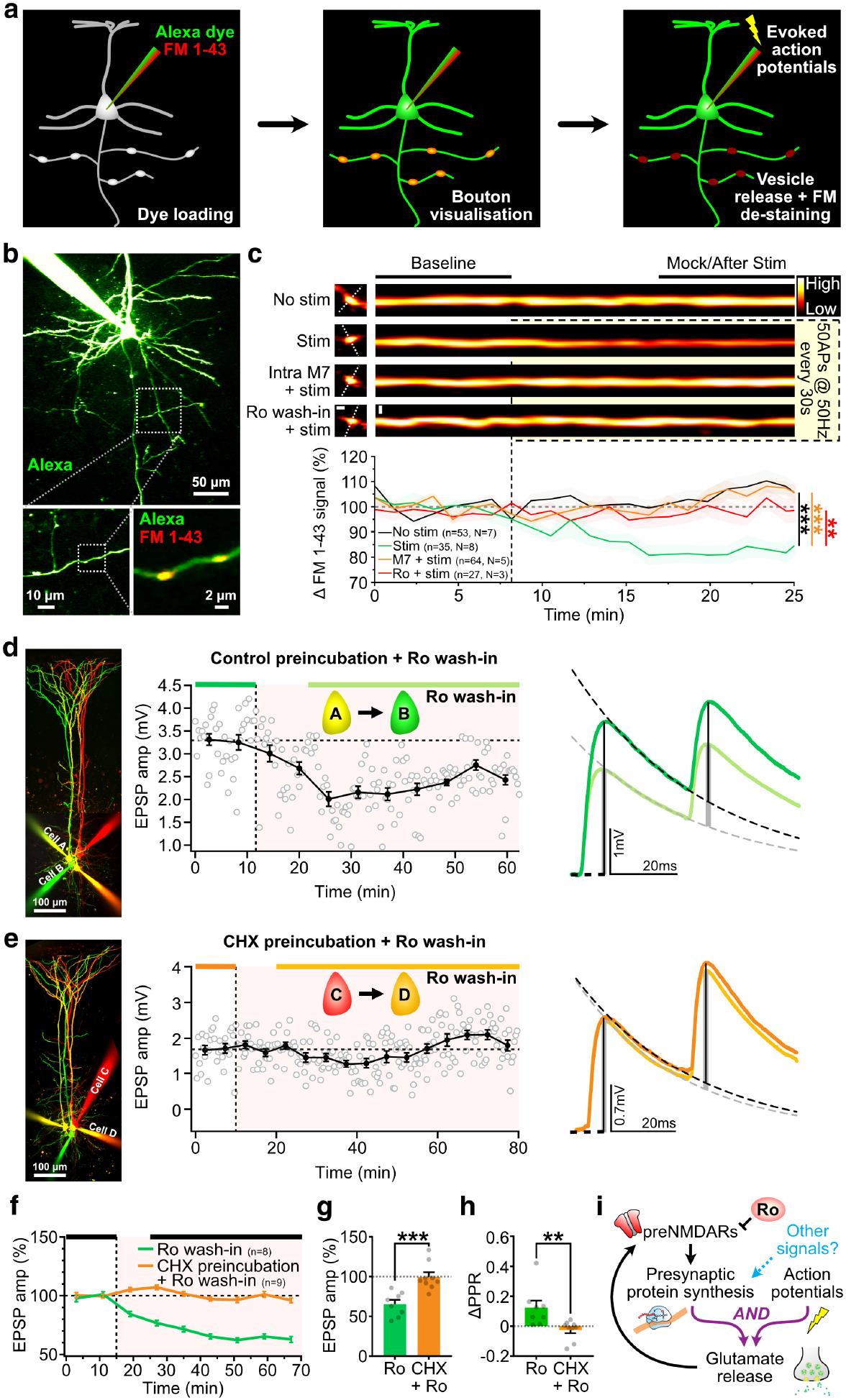
Presynaptic NMDARs sustain evoked vesicle release via translation. **a,** Neurons were loaded via the patch pipette with Alexa 488 and FM 1-43 dyes for visualization of morphology and of vesicle cycling, respectively. Evoked vesicle release led to FM 1-43 de-staining of presynaptic terminals. **b,** 2PLSM images of sample layer-5 PC loaded with vesicle dye FM 1-43 and morphological dye Alexa 488. Dotted boxes were enlarged to focus on axon and boutons. **c,** Top, FM 1-43 signal in sample boutons. Scale bars, 1 μm. Bottom, ensemble averages of FM 1-43 signal changes (*n = boutons, N = slices*). Inhibition of presynaptic translation by M7 loading abolished FM 1-43 de-staining and vesicle cycling, as did preNMDAR blockade with Ro 25-6981 (Kruskal-Wallis test, *P* < 0.001, \*\**P* < 0.01, \*\*\**P* < 0.001 with Dunn’s correction). Mean ± s.e.m. **d,** In controls, EPSPs were suppressed by Ro wash-in. Traces are averaged from baseline and after periods as indicated by color-coded lines. **e,** Ro-mediated EPSP suppression was abolished by CHX. **f,** Ensemble time courses show robustness. (*n = connections*). **g,** CHX preincubation occluded Ro-mediated EPSP suppression (*t_15_ = 4.35*). **h,** CHX occluded Ro-mediated PPR increase (*t_15_ = 2.97*). **I,** Schematic of tonically activated preNMDARs sustaining presynaptic protein synthesis and glutamate release.

Presynaptic *N*-methyl-D-aspartate receptors (preNMDARs) have been shown to sustain evoked PC→PC release by regulating the replenishment of the readily releasable pool of vesicles^33–35^. In agreement, wash in of Ro 25-6981 (Ro), which blocks GluN2B-containing preNMDARs at this developmental stage^33–35^, abolished FM 1-43 de-staining (Fig. 2c) and reduced EPSPs (Fig. 2d, f) in a manner similar to protein synthesis blockade (Figs. 1, 2c). We therefore explored if preNMDARs act via axonal protein synthesis by preincubating brain slices with CHX to inhibit translation (104 ± 7 min, n = 9 connections), followed by Ro wash-in. We found that translation inhibition occluded Ro-mediated changes in EPSPs, PPR and 1/CV^2^ (Fig. 2d-h and fig. S4b). These results suggest that preNMDARs can act upstream of presynaptic protein synthesis to sustain PC→PC release (Fig. 2i).

Having localized the need for protein synthesis to the presynaptic neuron, we next wondered whether it was local to the axon. To sever the presynaptic axon from the soma, we refined a 2-photon laser axotomy approach that we previously used^36^. In connected PC→PC pairs, we drove presynaptic spiking with an extracellular stimulation electrode placed close to the axon hillock (figs. S5, S6). This allowed us to sever the presynaptic soma from the axon with 2-photon laser axotomy (fig. S7) without cutting nearby neurites (fig. S8) and while still activating the connection of the PC→PC pair (Fig. 3a-c).

**Fig. 3.**
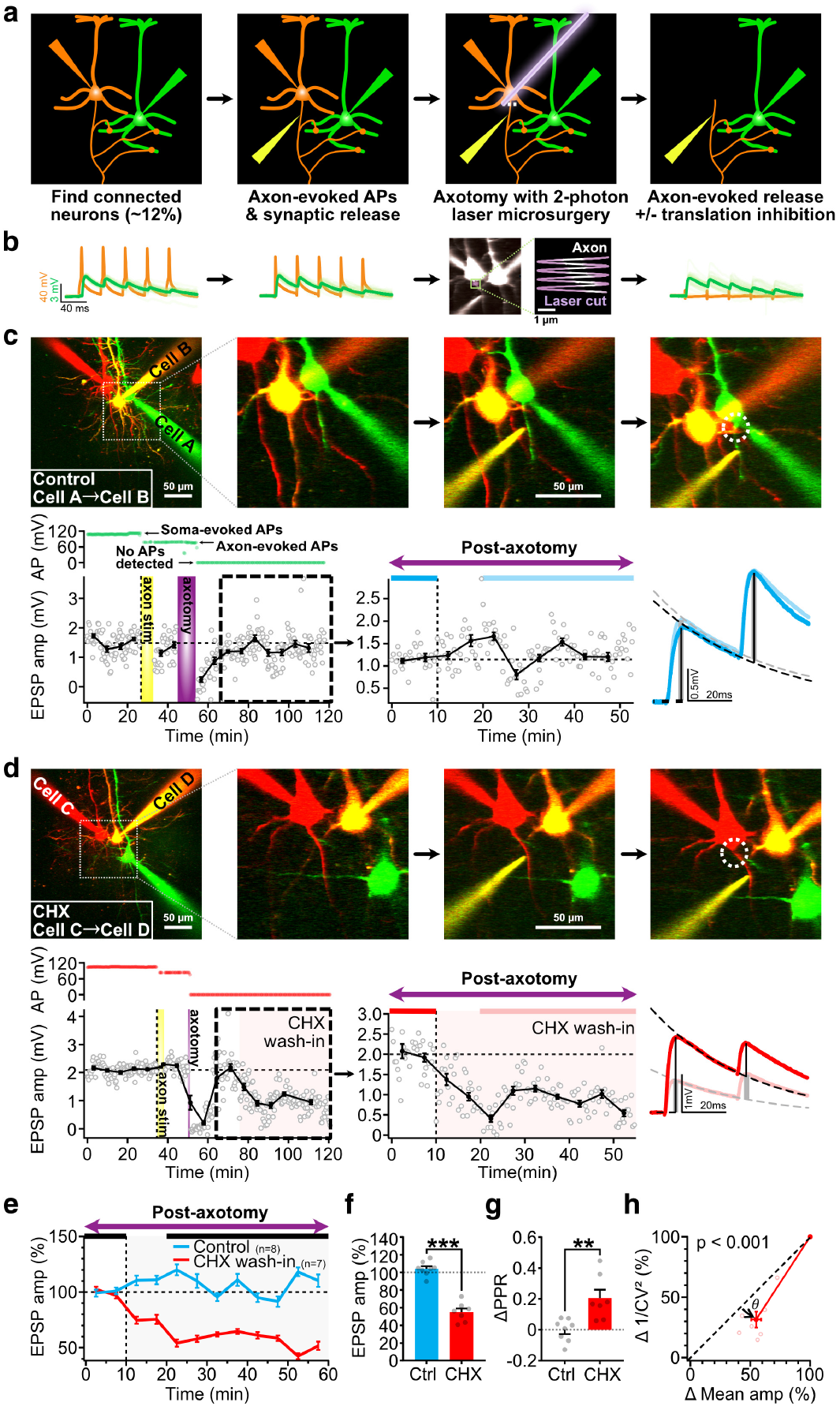
Local translation in axons supports evoked neurotransmitter release. **a,** Axonal stimulation (yellow) after 2-photon laser axotomy enabled testing the specific role of axonal protein synthesis in evoked release. **b,** Left, soma-evoked APs (orange) and EPSPs (green). Middle and right: axon-evoked APs and EPSPs before and after axotomy. Images showing the 4 × 2 μm sine-square scan (purple) used to cut the axon. Scale bar, 1 μm. **c,** Control experiment shows establishment of axon-evoked APs and corresponding EPSPs, followed by no presynaptic APs after axotomy, yet presence of EPSPs due to spiking isolated axon. Blue traces averaged from blue periods show stable synaptic responses. Scale bars, 50 μm. **d,** Within minutes, CHX reduced EPSPs (red traces and periods) elicited by stimulation of isolated axon. **e,** EPSP weakened post-axotomy with but not without CHX (*n = connections*) revealed that axonal but not somatic protein synthesis affected release. **f,** CHX reduced post-axotomy EPSPs compared to controls (*t_13_ = 10.0*). **g, h,** Analysis of PPR (*t_13_ = 3.54*) and CV (*Wilcoxon test, θ = 11° ± 2°, P < 0.001*) agreed that CHX reduced release. Mean ± s.e.m. Student’s t-test for (**f, g**). \*\**P* < 0.01, \*\*\**P* < 0.001.

We found that evoked release by isolated axons was sustained in the absence of the soma for more than an hour post axotomy (Fig. 3c). This suggests a high degree of axon autonomy in neurotransmission. In contrast, CHX wash-in reduced EPSPs in keeping with reduced evoked release from isolated axons (Fig. 3d-h). These results were indistinguishable from those obtained when axons were not cut (Fig 1 and figs. S1, S2, S3). Taken together, our findings strongly argue that translation specifically in the axon is necessary to sustain PC→PC release.

In agreement with axonal translation sustaining release, we observed a high abundance of endogenous RNA granules trafficking in main axons of PCs (Fig. 4a, b, Supplementary Video 1) like we previously saw in retinal axons *in vivo*^19^. In hippocampal dendrites, docked mRNA can be translationally repressed and undergo activity-dependent unmasking^37^. We hypothesized that synapses in neocortical PC axon collaterals are also capable of recruiting RNA. In agreement, we found that axonal boutons were rich in docked RNA granules (Fig. 4c; 168 out of 180 boutons; 93%, n = 7 cells), which is consistent with studies showing that ~80% of presynaptic terminals contain translation machinery^24,25^. These granules appeared to be essentially stationary when visually tracked to over an hour (Fig. 4b, d, Supplementary Video 2, 3), in stark contrast to the highly mobile granules found in main axons (Supplementary Video 1) and dendrites (Supplementary Video 4). To independently quantify RNA movement, we relied on a deep-learning approach^38^, which revealed a main axon granule speed of 14.4 ± 0.7 μm/min (Fig. 4e), similar to that reported in retinal axons^19,39^. However, granule speed in axon collaterals was 0.20 ± 0.07 μm/min, meaning RNA granules were mostly stationary. Since PC→PC synapses are formed by axon collaterals^40^, this finding is consistent with local axonal translation supporting neurotransmission.

**Fig. 4.**
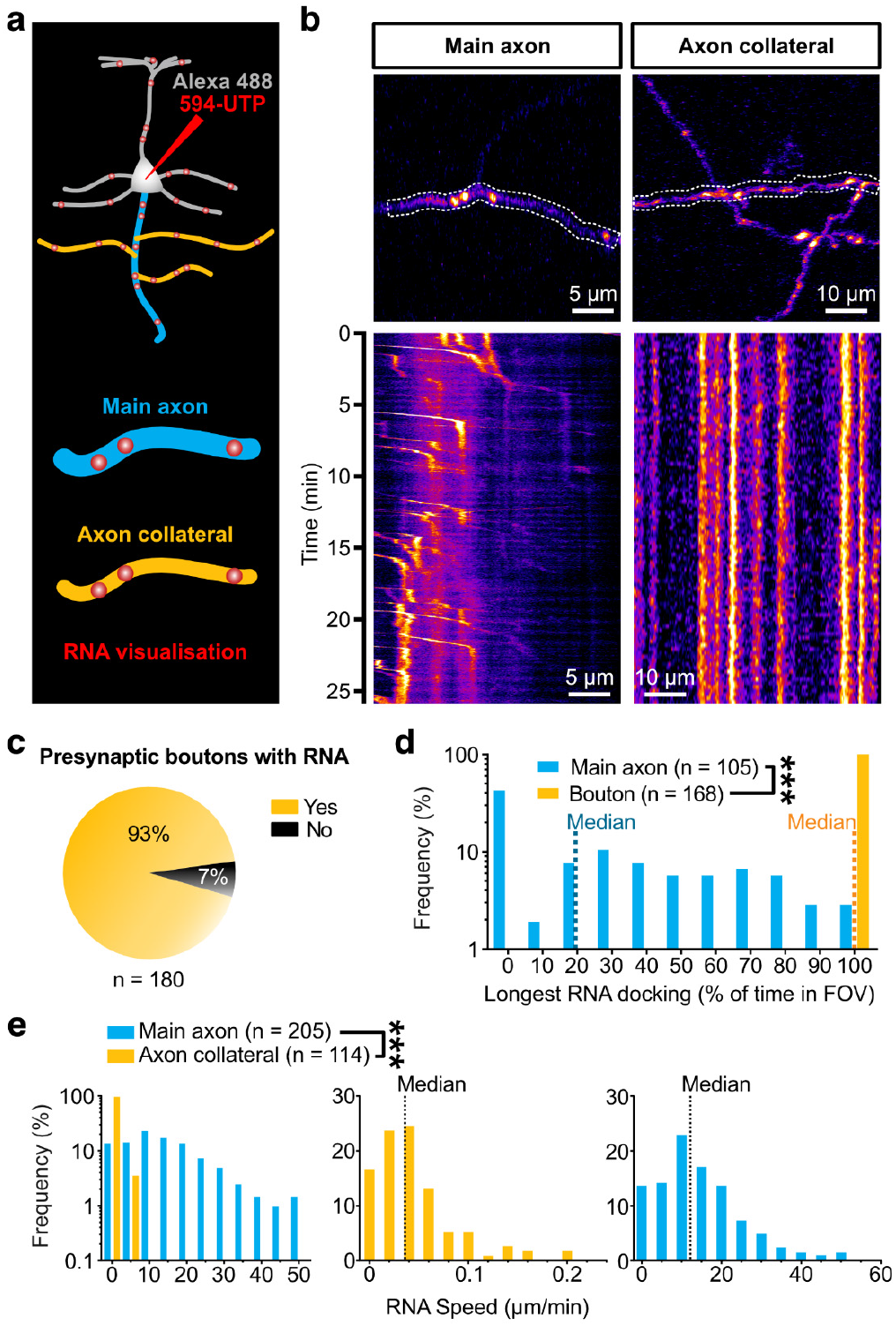
Endogenous RNA granules are stably docked at presynaptic terminals. **a,** Neurons were loaded via the patch pipette with Alexa 488 dye and Alexa 594-tagged uridine-5’-triphosphate (594-UTP) for visualization of cell morphology and endogenous RNA, respectively. **b,** Top: 2PLSM sample images of RNA granules in layer-5 PC main axon and axon collaterals, with soma located to the right. Bottom: Dotted boxes are plotted as kymographs. **c,** Proportion of boutons contained RNA granules (*n = 180 boutons, N = 7 cells*). **d,** Normalised longest RNA docking-time distribution show that RNA granules in axon collaterals (*n = 168 granules, N = 7 cells*) had longer docking time than those in the main axon (*n = 105 granules, N = 7 cells*). Dotted lines represent medians. **e,** Left: RNA granule speed distribution indicated lower speed in axon collaterals (*n = 114 tracks, N = 5 cells*) than in the main axon (*n = 205 tracks, N = 5 cells*). Middle and right: RNA speed distributions for axon collaterals and main axon. Dotted lines represent medians. Mann-Whitney test. *\*\*\*P* < 0.001.

We wondered if RNA granules docked in axon collaterals allow proteins to be locally synthesized for specific boutons, in which case protein synthesis blockade should specifically affect some but not all synapse types. We revisited our CHX wash-in experiments (fig. S1) and found occasional late-onset inhibitory responses (fig. S9a) characteristic of frequencydependent disynaptic inhibition (FDDI) mediated by Martinotti cells (MCs)^33,41,42^. Interestingly, this FDDI appeared to be resistant to inhibition of protein synthesis (fig. S9a).

Since we reasoned that axonal protein synthesis may regulate synapse types differentially, we thus wanted to explore EPSPs and FDDI simultaneously. To recruit FDDI effectively, we switched to a well-established high-frequency firing protocol^33,41,42^ (fig. S9b). We verified that presynaptic M7 suppressed PC→PC release equally well in this protocol (fig. S10). With this approach, we found that presynaptic M7 suppressed PC→PC but not PC→MC→PC connections (Fig. 5a-f), even for EPSPs and FDDI elicited by the same presynaptic PC (Fig. 5a-c). In separate experiments, we verified that protein synthesis did not affect FDDI by loading pre- or postsynaptic cells with CHX (fig. S9c). Taken together, our results argue that local translation in axons helps to sustain release at PC→PC (Figs. 1–3, figs. S2–4, S9a) but not at PC→MC→PC connections (Fig. 5 and fig. S9).

**Fig. 5.**
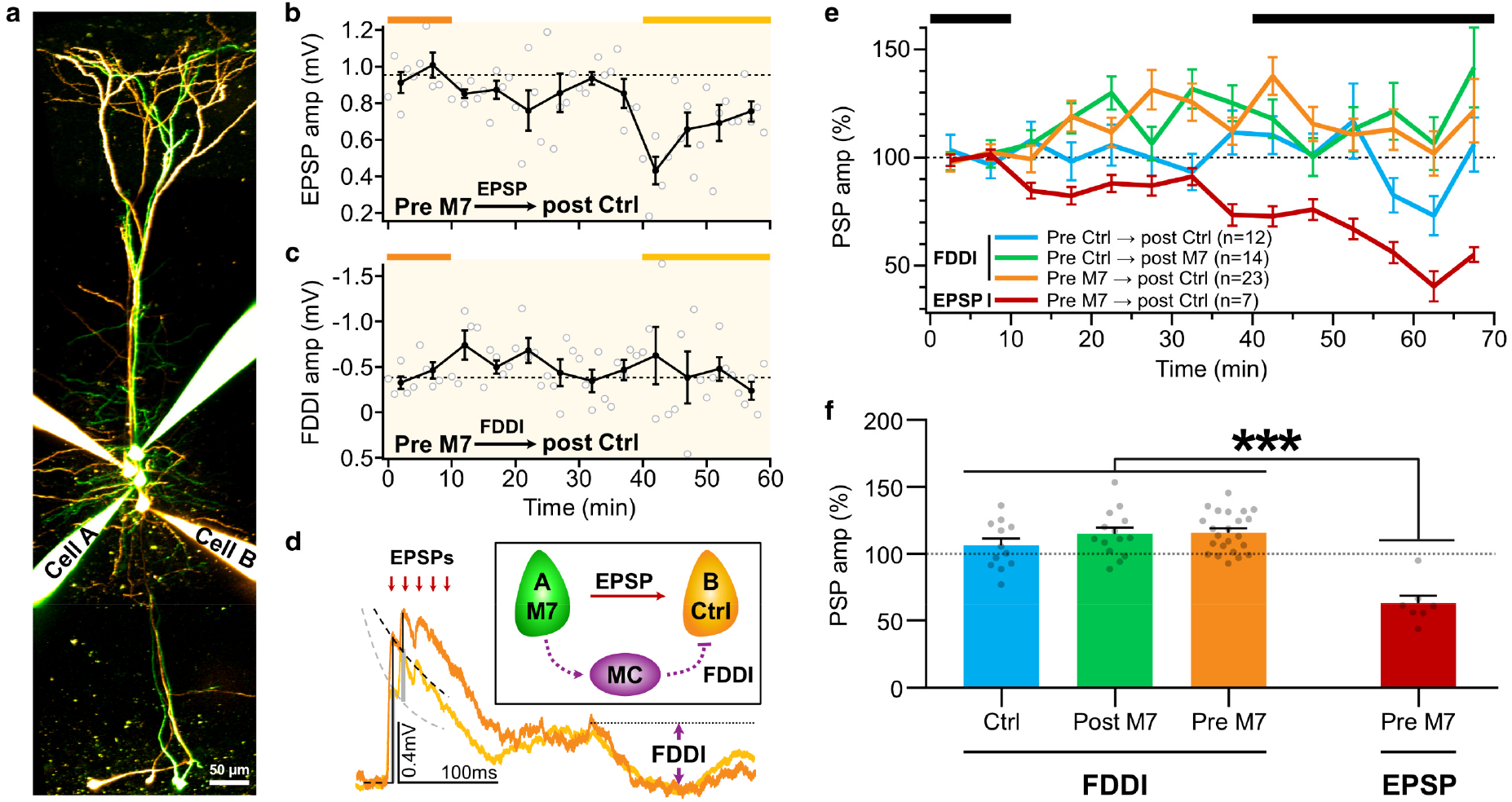
Presynaptic translation supports activity-dependent neurotransmission in a synapse-type-specific manner. **a,** 2PLSM image of a sample quadruple whole-cell recordings showing simultaneous monosynaptic excitatory connection and disynaptic inhibitory connection between Cell A (M7 loaded) and Cell B (control). **b,** EPSPs in monosynaptic connection with presynaptic M7 (Cell A). **c,** Frequency-dependent disynaptic inhibition (FDDI) with the same M7-loaded presynaptic cell. **d,** Traces are averaged from and color-coded by the baseline and “after” periods as indicated by bars above line graphs in (**b, c**). Inset displays the connectivity scheme, where Cell A (M7 loaded) forms a monosynaptic excitatory connection with Cell B. Cell A additionally forms a disynaptic connection with Cell B via a Martinotti cell (MC), giving rise to a delayed inhibitory component in the traces (purple arrow). **e,** Ensemble averages of PSPs (postsynaptic potentials) (*n = connections*). **f,** EPSP, but not FDDI, amplitude was suppressed by presynaptic translation inhibition (one-way ANOVA, *F_3, 52_ = 20.7, P* < 0.001; \*\*\**P* < 0.001 with Tukey correction). Mean ± s.e.m.

## Discussion

Our findings demonstrate that in the mammalian neocortex, protein synthesis in mature axons sustains evoked release at PC→PC connections via the mTOR and cap-dependent translation pathway. Protein synthesis, however, does not regulate PC→MC→PC synapses. Our study thus supports a model in which axonal mRNA translation occurs locally to differentially control specific synapse types^7,8^.

### The roles of axonal translation vary with development

Local translation provides an attractive explanatory framework for how neurons regulate functions far from the soma. Conventional genetic manipulations, however, lack the spatiotemporal precision to acutely and selectively manipulate translation in dendrites versus axons. Altered gene expression may thus not directly address specific local roles^4^. Even so, despite early scepticism, a consensus has emerged that local axonal translation supports axonal functions in early development^4,5,18^, such as β-actin in axon steering^43^ and branching^19,21^. Formation of presynaptic terminals in hippocampal cultured neurons also relies on axonal translation of β-catenin^23^ and SNAP25^22^ mRNA. Nonetheless, mRNAs encoding β-actin, β-catenin, and SNAP25 are not enriched in adult axons^24^, illustrating how axonal translation is regulated with development^26^.

Yet the role of local axonal translation in adult circuits is poorly explored. A pioneering study showed that in cultured neurons from juvenile *Aplysia* sea slugs, local protein synthesis determines branch-specific long-term plasticity^44^. These neurites, however, have juxtaposed pre- and postsynaptic domains, with unclear implications for local translation in mature mammalian axons and dendrites.

Two recent studies have shown that in mature hippocampus, long-term plasticity requires presynaptic translation, putatively in axons^25,27^. Our study, however, reveal that in mature axons of the neocortex, specific synapse types engage axonal translation to sustain release. Whether neurotransmission at specific synapse types in mature hippocampus depends on axonal translation remains unclear^1,2,25,27^. In general, however, these studies^25,27^ support our findings that local axonal translation subserves critical functions even in mature axons. To our knowledge, our study provides the first evidence that translation in mature axons can sustain information transfer at central synapses.

### Synapse-type-specific regulation of axonal translation

Here, we found that sustained neurotransmission at PC→PC synapses needed local axonal translation, suggesting that synaptic release requires efficient replenishment of key proteins. Responses mediated by PC→MC→PC synapses, however, were strikingly resistant to protein synthesis blockade. In agreement, translationally active boutons are known to vary with excitatory and inhibitory synapse types^24^. Consistent with synapse-type-specific axonal translation, we also found that RNA granules docked at a subset of axonal compartments, perhaps close to boutons requiring local axonal translation.

A distinct function of MCs is to mediate late-onset feedback inhibition of PCs^41,42^. This property results from PC→MC synapses that both strongly facilitate^41,42^ and release asynchronously^45^. However, PC→PC connections store information by recruiting long-term plasticity that is acutely sensitive to millisecond timing of pre- and postsynaptic spiking^46,47^. The differential regulation of PC→PC and PC→MC synapses thus likely reflects the distinct functions and needs of these connection types^7,8^.

### Mechanisms of mRNA regulation in axons

The differential requirement for protein synthesis of PC→PC and PC→MC synapses need not be static, however, but may instead be regulated differently by cues such as activity and neurotransmitters. For instance, we found that preNMDARs were linked to local axonal translation and regulated release at PC→PC connections. Although we identified preNMDARs as one candidate trigger, many other triggers may also initiate local axonal translation, such as the endocannabinoid CB1 receptor in hippocampal long-term depression of inhibition^25^.

RNA targeting permits local synthesis of essential synaptic proteins. For example, induction of chemical long-term potentiation was recently shown to enrich key mRNAs close to dendritic spines^48^. Furthermore, glutamate uncaging at spines enriched mRNA and locally synthesized β-actin for spine remodelling in an NMDAR-dependent manner^49^. It is therefore likely that preNMDARs may similarly recruit mRNAs to presynaptic boutons.

With over a thousand mRNAs in the axons^17,24,26^, how are specificity and timing of local translation achieved? One mechanism likely involves RNA-binding proteins — such as the fragile X mental retardation protein FMRP and the zip-code binding protein ZBP — which can translationally repress bound mRNA until use^50^. Another complementary model involves coupling receptors and translation machinery. Ligand binding may then enable local translation by releasing receptor-sequestered mRNA, ribosomes, and translation factors^51,52^. Local translation of specific mRNA subsets may thereby be triggered in a ligand-specific^53^ or learning-associated manner^17^. Since NMDAR activation has been shown to unmask^37^ and activate mRNA translation in dendrites^37,48,49^ and since we link preNMDARs to translation-dependent release, it will be interesting to investigate whether NMDARs have a dedicated mRNA interactome^51,52^.

### Future Directions

Recent studies have revealed how specific neuronal populations engage *de-novo* protein synthesis to mediate learning and behaviour^54–56^. Indeed, correcting protein synthesis is emerging as a therapeutic target for neuropathology ranging from autism to Alzheimer^5,57^. One proposed strategy has been to modulate protein synthesis to restore excitatory/inhibitory balance^58^, which is often derailed in autism spectrum disorder^58,59^. For example, protein synthesis and excitation are elevated in fragile X syndrome, although most studies focus on postsynaptic changes^58^. One study, however, showed that axons in the adult human brain contain the fragile X translational regulator FMRP^60^. Our findings reveal how presynaptic protein synthesis in mature axons boosts excitatory PCs but not inhibitory MCs, highlighting the prospects of recalibrating neocortical excitatory/inhibitory balance in disease by targeting local presynaptic translation.

By combining quadruple patch clamp, live imaging, and laser microsurgery, we uncovered how local axonal translation sustains neurotransmission at specific neocortical synapse types. Local axonal protein synthesis is therefore a plausible determinant of synapsetype-specific plasticity^7,8^. In conjunction with other powerful techniques^24,26,53,61^, our approach will thus reveal the diversity of local synaptic regulation in neural circuits.

## Methods

All animal procedures conformed to the Canadian Council on Animal Care as overseen by the Montreal General Hospital Facility Animal Care Committee, with appropriate licenses.

### Slice Preparation and Basic Electrophysiology

C57BL/6J mouse brains from either sex were dissected on postnatal day (P) 11-16 in ice-cold (<4°C) artificial cerebrospinal fluid (ACSF: 125 mM NaCl, 2.5 mM KCl, 1.25 mM NaH_2_PO_4_, 26 mM NaHCO_3_, 25mM D-Glucose, bubbled with 95% O_2_/5% CO_2_, osmolality adjusted to 338 mOsm.kg^−1^ with D-glucose). Using a Campden Instruments 5000mz-2 vibratome, brain slices of 300 μm were cut from the visual cortex at angle normal to the pial surface to maximally preserve apical dendrites and principal axons. Slices were transferred to incubation chamber at 32°C for 20 min before cooling down to room temperature (~23°C). Experiments were performed with ACSF perfused at 32-34°C, maintained by a resistive inline heater (Scientifica). Temperature was recorded and verified offline. Recordings were discarded or truncated if temperature deviated outside this range.

Patch pipettes of 4-6 MΩ resistance were pulled from standard-wall borosilicate capillaries (G150F-4, 1.5 mm outer diameter, 0.86 inner diameter, Harvard Apparatus) with a P-1000 electrode puller (Sutter Instruments). Pipettes were loaded with internal solution (5 mM KCl, 115 mM K-Gluconate, 10 mM HEPES pH 7.4 calibrated with KOH, 4 mM Mg-ATP, 0.3 mM Na-GTP, 10 mM Na-phosphocreatine, 0.1% w/v Biocytin, adjusted with KOH to pH 7.4, osmolality adjusted to 310 mOsm/kg with sucrose). For 2PLSM, internal solution was supplemented with 20-150 μM Alexa Fluor 488 and/or Alexa Fluor 594 (Life Technologies).

Whole-cell recordings were performed using MultiClamp 700B amplifiers (Molecular Devices). Current clamp recordings were filtered at 20 kHz and acquired at 40 kHz using PCI-6229 boards (National Instruments) with custom-written software^62^ (https://github.com/pj-sjostrom/MultiPatch.git) running in Igor Pro 7, 8 or 9 (WaveMetrics). Series resistance, input resistance, resting membrane potential or holding current, excitatory postsynaptic potential (EPSP) and perfusion temperature were monitored online and assessed offline. Series resistance was continuously monitored but not compensated. Recordings were not adjusted for liquid junction potential (10 mV).

Neurons were patched at 400× magnification with infra-red video Dodt contrast on a custom-modified microscope based on the BX51WI frame (Olympus). Primary visual cortex was identified based on the presence of layer 4. Layer-5 PCs were identified based on their characteristically large pyramid-shaped somata and thick apical dendrites. Neuronal morphology was verified *post hoc* using 2PLSM of Alexa fluorescence.

### Two-photon Laser Scanning Microscopy (2PLSM)

2PLSM was performed with an imaging workstation custom-built from a BX51WI microscope (Olympus). Detectors were based on the R3896 bialkali photomultiplier tubes (PMT; Hamamatsu). Scanners were 6215H 3-mm galvanometric mirrors (Cambridge Technology). Two-photon excitation was achieved using a MaiTai HP titanium-sapphire laser (Spectra-Physics) tuned to 750-840 nm. Lasers were gated with SH05/SC10 (Thorlabs) shutter-controller pair, and manually attenuated with a polarizing beam splitter in combination with a half-wave plate (Thorlabs GL10-B and AHWP05M-980). Laser power output was monitored with a power meter (Thorlabs PM100A with S121C). Fluorescence was collected with a FF665-Di01 or FF695-Di01 dichroic (Semrock). Green and Red fluorescence were separated with a FF560-Di01 dichroic beam mirror (Semrock), a ET525/50m (Chroma) green emitter, and ET630/75m (Chroma) red emitter. Laser-scanning Dodt contrast was obtained by collecting the laser light after the spatial filter with an amplified diode (Thorlabs PDA100A-EC). Imaging data were acquired using ScanImage^63^ versions 3.7 (customized), 2019, 2020 or 2021 running in MATLAB (The MathWorks) via PCI-6110 or PCIe-6374 data acquisition boards (National Instruments).

After quadruple recordings, layer-5 PC morphologies were acquired as stacks of 512×512 or 1024×1024-pixel image slices separated by 1-2 μm. Each slice was an average of three acquisitions at 2-4 ms per line. Four stacks were typically needed to cover the whole dendritic and axonal trees for each PC.

### Pharmacology

Cycloheximide (CHX; Bioshop Canada # CYC003) was washed-in at 80 μM. Ro 25-6981 maleate (Ro; Hello Bio #HB0554) was washed-in at 0.5 μM. For intracellular loading via patch pipettes, 80 μM CHX, 5 μM PP242 (Tocris #4257) or 120 μM m7G(5’)ppp(5’)G RNA Cap Structure Analog (M7; New England BioLabs #S1404S) was used. For PP242, DMSO was used as a solvent and hence an equal concentration (1:5000) of DMSO was added into the internal solution for patching control cells.

### Puromycin-Based Translation Assay

Assay was adopted from previously reported *in-vitro*^64^ and *in-vivo*^19^ approaches with the use of puromycylation of nascent proteins and immunolabelling of puromycin. Brain slices were first incubated in control ACSF, 40 μM, 80 μM or 120 μM CHX for 10 min at 32°C, followed by incubation with 10 μg/ml puromycin (Sigma) for 10 min, except for Puro Nil controls. Slices were next washed with ACSF for 5 min to remove unincorporated puromycin prior to fixation with 4% paraformaldehyde in 1× phosphate-buffered saline (PBS: 10 mM NaCl, 2.7 mM KCl, 10 mM NaH_2_PO_4_, 1.8 mM KH_2_PO_4_) at 4°C overnight. Fixed slices were washed 5 times with 1 × PBS + 1% Triton X-100 (PBST) for 10 min each before incubating with 4% heat-inactivated goat serum in PBST at 4°C overnight. Alexa Fluor 488 conjugated anti-puromycin clone 12D10 antibody (Millipore MABE343-AF488; 1:100) was incubated with slices in PBST at 4°C for 2 days. Immunostained slices were washed 5 times with PBST for 10 min and twice with 1× PBS for 5 min to remove detergent. They were then mounted on slides with ProLong Gold Antifade Mountant (Life Technologies) and protected with coverslips.

2PLSM was employed to acquire images from mounted slices at 780 nm, with the laser power and photomultiplier gain being kept the same for all conditions in each experiment. Each 60-μm-thick image stack at 5 μm intervals was taken at 512×512 pixels and averaged from 8 acquisitions. Three to four fields of view (FOVs) were taken from each slice and three to four slices were used for each condition per experiment. The Puro Nil control was added in the last two out of three experiments, with four FOVs from each of the seven slices. Background for each FOV was measured by a 100×100-pixel square in an empty area of the top layer in the stack as an average. The average puromycin fluorescence for each FOV was measured after maximally projecting the stack, followed by subtracting the background. The background-corrected values were normalized to the mean of the control condition.

### Evoked Neurotransmission

Layer-5 PCs are sparsely connected (< 12%)^28^, so we compensated the experimental yield by using quadruple whole-cell recordings to test for 12 possible connections simultaneously^65^. Gigaohm seals were formed with four cells and then broken through in quick succession. To test for connections, 5 spikes were evoked at 30 Hz by 5-ms-long current injections of 1.3 nA every 20s for 10 repetitions. To ensure long-term plasticity was not unintentionally induced, spikes in different cells were separated by >700 ms^62^. If EPSPs were absent, all four recordings were discontinued, and the experiments were restarted with fresh pipettes. When at least one connection with sufficiently large EPSP (>0.3 mV) was found, baseline recording was commenced.

All evoked neurotransmission experiments were done in current clamp. FDDI experiments (Fig. 5 and figs. S9b, S9c, S10) were evoked with a burst of 15 spikes at 70 Hz every 60 s^33,41,42^. For the rest, bursts of 5 spikes at 30 Hz were evoked every 20s. Resting membrane potentials, input resistances, EPSP amplitudes and temperature were continuously monitored online. If these properties were stable, wash-in of CHX or Ro was commenced after 10-20 min of baseline recordings. For experiments with intracellular loading of pharmacological agents, minimal positive pressure was applied just before pipettes entered the slice.

Quality selection criteria similar to those previously established were used^33,34,62^. Input and series resistance were assessed with a 250-ms-long test pulse of −25 pA. Recordings with more than 30% change in input resistance or more than 8 mV change in resting membrane potential were discarded or truncated. Recordings shorter than 20 min after commencing drug wash-in were not used. The larger-size drug M7 may take longer to reach axon terminals as suggested by previous report^25^. Therefore, M7 experiments shorter than 40 min after the baseline period were not used. Experiments with unstable baseline were discarded, with stability measured using Student’s t-test of Pearson’s r for EPSP amplitude versus time.

Evoked responses were averaged over before and after periods, as indicated in figures. Suppression of evoked neurotransmission was presented as a ratio of the first response in a train averaged over drug wash-in or post-loading divided by that averaged over the pre-drug or pre-loading baseline period. Short-term plasticity was measured using the PPR, defined as (R_2_-R_1_)/R_1_, where R_i_ is the i^th^ response. Trains of 5 or 15 responses were acquired. However, as it was previously found that using R_3_ and beyond did not appreciably affect analysis^66^, we relied solely on PPR. The change in PPR (ΔPPR) was calculated as RRP_after_ -PPR_before_. CV analysis was performed as previously described^31^. Briefly, mean EPSP and CV of R_1_ were calculated for baseline and drug conditions, followed by normalizing mean EPSP and CV^−2^ to the baseline period. The angle (*θ*) was defined by the diagonal unity line and the line formed by linking the starting coordinate (100%, 100%) and CV analysis endpoint. Finding *θ* < 0 (i.e., clockwise from diagonal) indicated postsynaptic locus, while *θ* > 0 indicated presynaptic locus (Fig. 2G).

### FM 1-43 vesicle release visualization assay

To enable high signal-to-noise visualization of vesicle release at presynaptic boutons, we carried out FM 1-43 imaging of vesicle cycling^32^. We loaded 5 μM FM 1-43 together with 150 μM Alexa Fluor 488 morphological dye into layer-5 PCs for at least 1 hour with large-opening pipettes (2 – 4 MΩ). Burst of 5 spikes at 30 Hz were evoked every 20 s for 30 min to facilitate dye loading and then stopped thereafter to avoid de-staining. Boutons were identified using Alexa Fluor 488 at 3× to 5× zoom (103×103 to 62×62 μm FOV) and 256×256 pixels, with 2-photon laser tuned to 750 nm. Time-lapse imaging of FM 1-43 was performed using 256×256-pixel z-stacks with slices separated by 1 μm, acquired every 70 s at 5× zoom. The laser was tuned to 840 nm for optimal FM 1-43 excitation, with Alexa Fluor 488 being weakly excited for post-experiment morphological registration. Each slice was an average of three acquisitions at 2 ms per line. After a baseline of 8 timepoints (490 s), the PCs were stimulated to fire 50 action potentials at 50 Hz every 30 s apart from the no stim control condition.

To register FM 1-43 time-lapse images, image stacks were corrected for drift in x, y, and z. In FIJI^67^, a region of interest (ROI) of the axon and/or dendrite was selected in a single z-plane of the stack, followed by the use of the *Correct 3D drift* script^68^ for the morphological dye Alexa Fluor 488. For each bouton, the brightest plane was identified and then a maximum projection was created by also including two slices above and two slices below to ensure signal coverage. From the flattened image, a ROI covering the bouton was created with the oval tool and the integrated FM 1-43 signal was recorded across the time-lapse images. The same ROI was subsequently moved to a nearby background area and with signal also recorded across the time-lapse images. The FM 1-43 signal in the bouton was then background corrected. Experiments with unstable baseline were discarded, with stability measured using Student’s t-test of Pearson’s r for signal versus time.

### Axon stimulation

Layer-5 PCs were filled with Alexa dye and imaged with 2PLSM. Connections were identified (see above). The axons were identified based on their smooth, non-spiny appearance and their projection down to the white matter. The axon initial segment (AIS) was targeted ~10-20 μm along the axon from the soma. Stimulation pipettes (5 – 6 MΩ) were filled with ACSF supplemented with Alexa dye. Aided by 2PLSM and laser-scanning Dodt-contrast, stimulation pipettes were positioned ~1 μm from the AIS target. Care was taken not to damage the axon. Five 100-ms biphasic spikes each in a voltage ramp of 0 – 18 V in 2 V steps were applied via the stimulation pipettes with 1 s inter-ramp intervals (fig. S6a). This ramp was repeated 3 – 5 times to identify a consistent minimum stimulation voltage threshold required to elicit presynaptic spiking, measured as antidromic responses with the somatic pipette. The minimal stimulation that triggered axon-evoked APs simultaneously elicited EPSPs in the postsynaptic cell.

To verify the specificity and spatial tolerance of axon stimulation, a set of control experiments was performed (fig. S5). One spike each in a voltage ramp of 0 – 18 V in 2 V steps were applied via the stimulation pipettes with 1s inter-ramp intervals, and the sweep was repeated 5 times. The success rate of AP firing was tallied for each stimulation voltage (fig. S5b). This was repeated each time the stimulation pipette was retracted diagonally with a set revolution on the micromanipulator. To correct for physical resistance from the tissue during the retraction, the closest diagonal stim-axon distance, *c*, was calculated by the Pythagoras theorem, *c^2^ = a^2^ + b^2^* (fig. S5c). The xy distance from the pipette tip to the point on the same plane directly above the closest point of the axon defines *a*, while the z distance between the tip and axon defines *b*. The AP firing rate at minimum stimulation voltage threshold was then plotted against the stim-axon distance (fig. S5d). On average, the AP firing probability dropped sharply below 50% when the pipette moved ~1.2 μm away from the stimulation origin, indicating high specificity. In a separate analysis, the stimulation strength required for full AP firing fidelity was charted against the stim-axon distance (fig. S5e). The data revealed that 120% of minimum stimulation voltage provides an average spatial tolerance of ~1.2 μm from the stimulation origin. To avoid interrupting recordings by minor tissue movement and to ensure fidelity of axon-evoked APs, 120% of minimum stimulation voltage was employed in experiments.

### Two-photon laser microsurgery

Three protocols of laser axotomy microsurgery were tested for separating the axon from cell soma: 1) 10 μm line cut at 1000 Hz, 2) 4×5 μm (width × height) sine-square cut at 125 Hz, and 3) 4×2 μm sine-square cut at 125 Hz. A unit of cut was performed for 500 ms every 10 s, with laser tuned to 780 nm and a back aperture power of 50 mW. Each cut was externally triggered by Igor Pro that simultaneously began electrophysiology recordings. Successful axotomy attempts were faithfully typified by 1) the attenuation of antidromic responses, 2) rapid depolarization of presynaptic soma, and 3) a concomitant drop in input resistance (Fig. 3 and fig. S7). These characteristics were live-monitored, and an additional cut was performed if this signature was not found. This ensured only the minimally required laser microsurgery was delivered and facilitate the preservation of target axon and surround tissue (fig. S8). The efficiency of axotomy was compared for the three protocols and showed that they were similar in terms of total laser axotomy time. 2PLSM images showed that axon preservation was typically better with the 4×2 μm sine-square protocol than 4×5 sine-square protocol (e.g., fig. S8), and was therefore opted for experiments (Fig. 3). The laser microsurgery was efficient at least up to the depth tested (~100 μm; fig. S7b). Overall, a median of 2 s of total laser time was required, providing a rapid means to perform axotomy with μm precision.

### Endogenous RNA labelling and dynamics tracking

Endogenous RNA was labelled by the Alexa-594 UTP analogue (Jena Bioscience; NU-821-PEG5-AF594), similar to previous studies^19,39,69,70^. UTP analogues were confirmed to be incorporated selectively into RNA, including mRNA and ribosomal RNA, but not genomic or mitochondrial DNA^19^. To visualize arbour and boutons, 150 μM Alexa Fluor 488 morphological dye was loaded intracellularly into layer-5 PCs with 6.25 μM Alexa-594 UTP via patch pipettes. Burst of 5 spikes at 30 Hz were evoked every 20 s to facilitate loading. To allow UTP incorporation into RNA, 2PLSM live-imaging of RNA granules were commenced at least after 2 hours of loading. Boutons were identified using Alexa Fluor 488 at 3×-5× zoom (103×103 to 62×62 μm FOV) and 256×256 pixels, with 2-photon laser tuned to 780 nm. Time-lapse imaging of RNA granules in axon collaterals was performed using z-stacks with slices separated by 1 μm, acquired every 11-30s at 5× zoom. For the main axon, time-lapse imaging performed at a single z plane and acquired every at 1.48 Hz at 5× zoom. Each time-lapse was performed for at least 20 minutes, and up to over an hour for verification of docking stability of RNA in boutons.

To register time-lapse images, image stacks were corrected for drift in x, y, and z as described above for FM 1-43. Longest period of docking was quantified for RNA granules, with docking defined as stationary or oscillatory movements within the diameter of the granule^19^. This period was normalized to the time of the RNA granules stayed in the field of view. Independently, kymographs of RNA tracks were analysed with the automated and artificial intelligence-assisted software KymoButler^38^. Thresholds for track detection was set to the default values of 0.1 - 0.2. Kymographs along the main axon or axon collaterals were prepare with KymographBuilder (https://github.com/fiji/KymographBuilder/) and the segmented line tool in FIJI^67^.

### Statistical analysis

Data were analysed in Igor Pro 9 or PRISM 7 (GraphPad). Results are presented as mean ± standard error of mean (s.e.m.), and *n* represents the number of connections in multiple patch-clamp experiments and number of neurons in other experiments, unless stated otherwise. Data from both sexes were pooled. All statistical tests were two-sided. Two-sample comparisons were made using unpaired Student’s t-test. If the F test for comparing variances was significant (*P* < 0.05), the unequal-variances Welch’s t-test was used. For RNA docking time and speed, which did not distribute normally, the Mann-Whitney *U* test was used. When faced with multiple comparisons, two-sample comparisons were performed only if one-way ANOVA permitted it at the *P* < 0.05 level. Kruskal-Wallis test was used if D’Agostino & Pearson normality test showed non-normal distribution at the *P* < 0.05 level. Multiple comparisons were corrected *post hoc* with Tukey’s for one-way ANOVA and Dunn’s for Kruskal-Wallis test. When applicable, F values and t values are presented in figure captions. Correlation tests were performed with both Pearson’s r and Spearman’s rho. For CV analysis, Wilcoxon’s rank test was used to compare the angle *θ* to the diagonal, i.e., 45°. Significance levels are denoted as * for *P* < 0.05, ** for *P* < 0.01 and *** for *P* <0.001.

## Acknowledgments

We are grateful to Arkady Khoutorsky, Charles Bourque, and Keith Murai for critical reading of the manuscript. We thank members of the Sjöström lab for help and useful discussions. H.H.-W.W. was supported by CIHR Fellowship 295104, HBHL Postdoctoral Fellowship, FRQS Postdoctoral Fellowship 259572 and QBIN Scholarship 35450. A.J.W. received funding from NSERC Discovery grant 2016-05118, CIHR PGs 153150 and 173290. P.J.S. acknowledges funding from CFI LOF 28331, CIHR PG 156223, FRSQ CB 254033, and NSERC DG/DAS 2017-04730.

## Author contributions

H.H.-W.W. and P.J.S. conceived of the project and of the methodology. H.H.-W.W. performed experiments and analysed data. P.J.S. supervised the project and wrote custom data acquisition and analysis software. H.H.-W.W., A.J.W., and P.J.S. wrote the manuscript.

## Data availability

The datasets generated during and/or analysed during the current study are available from the corresponding author on reasonable request.

## Code availability

All custom code used for data acquisition and data analysis are available from the corresponding author on reasonable request, as well as in GitHub links provided in the Methods section.

## Competing interests

Authors declare that they have no competing interests.

**Correspondence and requests for materials** should be addressed to H.H.-W.W. or P.J.S.

---

**Supplementary Video 1 | RNA granules are dynamically trafficked in the main axon**

2PLSM time-lapse imaging of layer-5 PC axon shaft and RNA granules.

**Supplementary Video 2 | RNA granules are stably docked in axon collaterals for tens of minutes**

2PLSM time-lapse imaging of layer-5 PC axon collaterals and RNA granules.

**Supplementary Video 3 | RNA granules are stably docked in axon collaterals for > 1 hour**

2PLSM time-lapse imaging of layer-5 PC axon collaterals and RNA granules.

**Supplementary Video 4 | RNA granules trafficking in dendrite but not in axon collaterals**

2PLSM time-lapse imaging of layer-5 PC basal dendrite, axon collaterals and RNA granules.

**Fig. S1.**
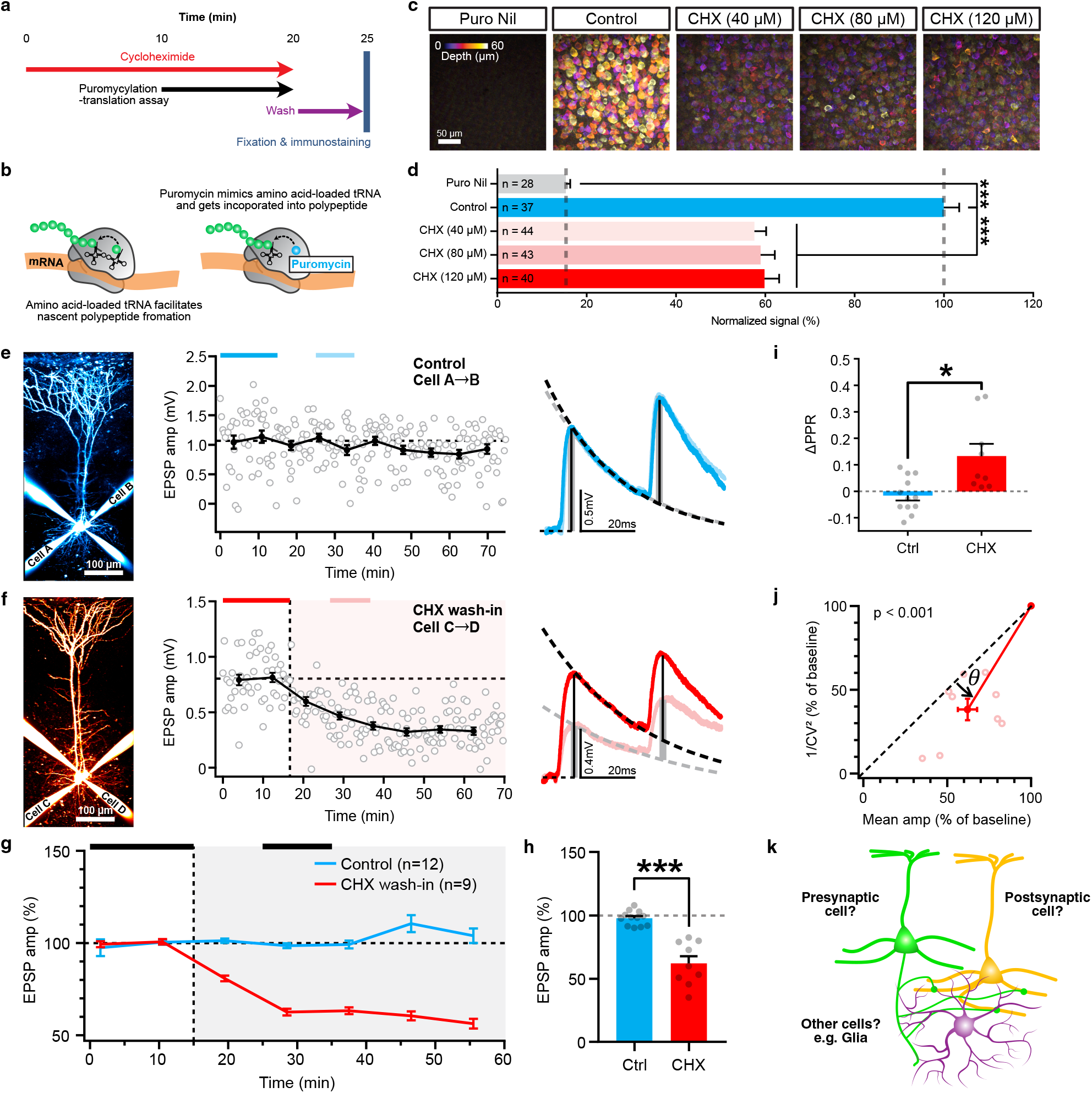
Protein synthesis inhibition suppresses evoked release. **a,** Timeline of translation assay post-cycloheximide (CHX) incubation. **b,** Translation elongation makes use of aminoacyl tRNA to synthesize polypeptide. Puromycin, an aminonucleoside antibiotic, is a structural analogue of aminoacyl tRNA and is incorporated into the nascent polypeptide chain. This can be exploited as an assay of translation efficiency with the use of antibodies against puromycin for visualization**^19,22,24,53–56,64,71^**. **c,** Images of puromycin incorporation color-coded by depth. Scale bar, 50 μm. **d,** Quantification of puromycin incorporation (Puro nil: n = 28 field of view (FOV), 7 slices; Control: n = 37 FOV, 10 slices; 40 μM CHX: n = 44 FOV, 11 slices; 80 μM CHX: n = 43 FOV, 11 slices; 120 μM CHX: n = 40 FOV, 10 slices; Kruskal-Wallis test, *P* < 0.001, \*\*\**P* < 0.001 with Dunn’s correction). **e,** Sample quadruple whole-cell recordings showing Cell A is connected to Cell B in the control condition. EPSPs remain stable over the course of over 70 min. Traces are averaged from and color-coded by the baseline and “after” periods as indicated by bars above the line graph. **f,** Same as (**e**) for weakened EPSPs in the connection from Cell C to Cell D in the CHX wash-in condition. **g,** Ensemble averages. (n = connections). **h,** CHX wash-in for 10 min reduced EPSP amplitude. Statistics are shown for 10 to 20 min after wash-in to illustrate the fast action after CHX wash-in (Control vs. CHX: 97.7 % ± 2 % vs. 62.1 % ± 6 %, *t_9.6_ = 5.99*, unequal variance t test, \*\*\**P* < 0.001). Similar statistics are obtained when the full time-courses are included (Control vs. CHX: 99.2 % ± 5 % vs. 59.8 % ± 6 %, *t_19_ = 5.15*, student t test, *P* < 0.001). **i,** CHX washin increased PPR, indicating short-term facilitation (*t_10.9_ = 2.99*, unequal variance t test, \**P* < 0.05). **j,** CV analysis showing points are below the diagonal for connections with presynaptic M7 loading (Wilcoxon test, *θ* = 14° ± 2°, *P* < 0.001), indicating a presynaptic locus of depression. **k**, The locus of translation inhibition that gives rise to suppressed neurotransmission cannot be differentiated with CHX wash-in. Mean ± s.e.m.

**Fig. S2.**
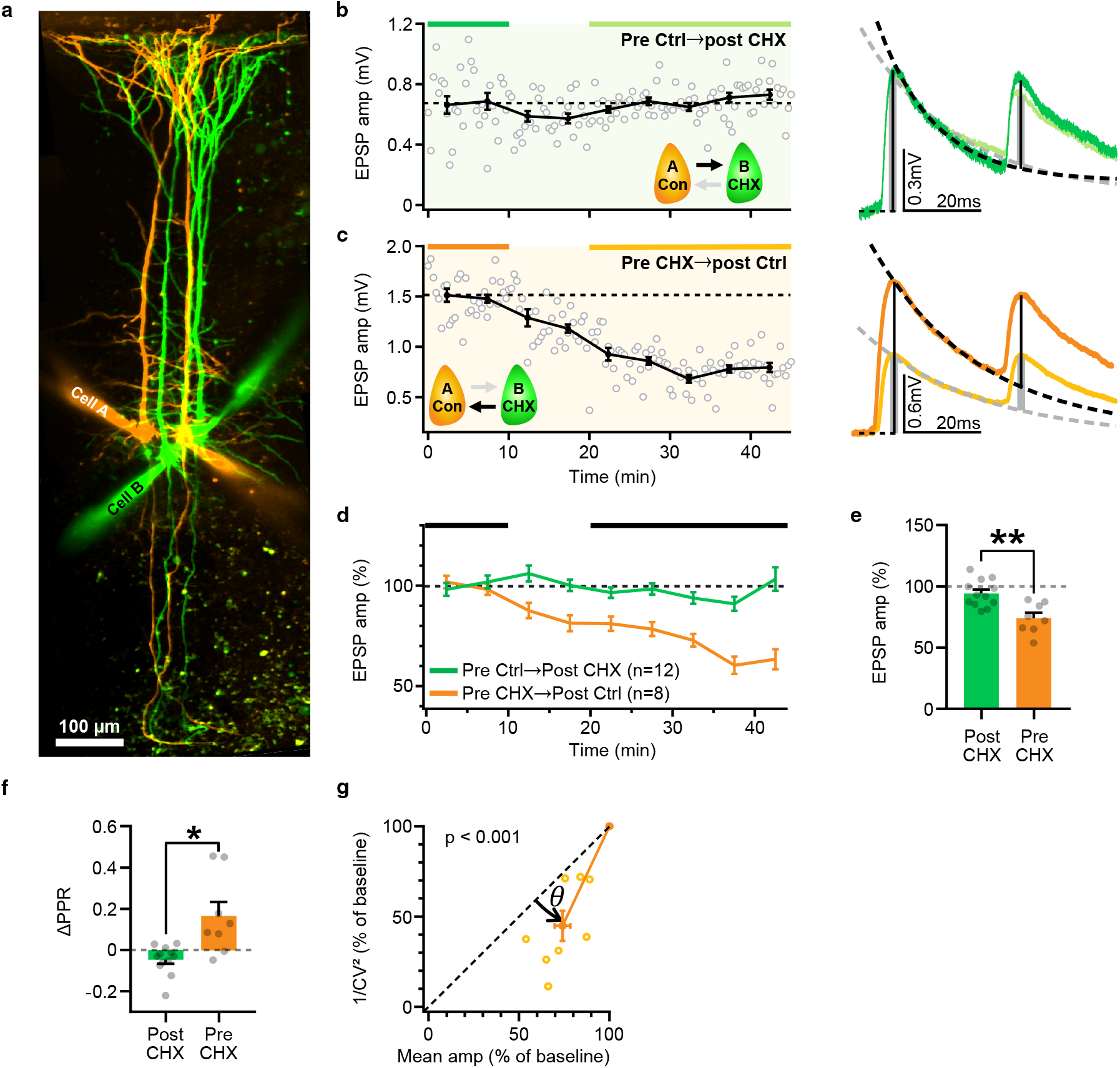
Presynaptic translation supports evoked release. **a,** 2PLSM image of a sample quadruple whole-cell recordings showing bidirectional connections between Cell A (control) and Cell B (CHX-loaded). Scale bar, 100 μm. **b,** Stable EPSPs in connection with postsynaptic CHX (Cell B). Traces are averaged from and color-coded by the baseline and “after” periods as indicated by bars above the line graph. **c,** Same as (**b**) for weakened EPSPs in the reciprocal connection with presynaptic CHX (Cell B). **d,** Ensemble averages. (n = connections). **e,** Presynaptic, but not postsynaptic, CHX loading reduced EPSP amplitude (*t_18_ = 3.84*, unpaired t test, *\*\*P* < 0.01). **f,** Presynaptic CHX loading increased PPR, indicating short-term facilitation (*t_8.3_ = 2.99*, unequal variance t test, \**P* < 0.05). **g,** CV analysis showing points are below the diagonal for connections with presynaptic M7 loading (Wilcoxon test, *θ* = 19° ± 3°, *P* < 0.001), indicating a presynaptic locus of depression. Mean ± s.e.m.

**Fig. S3.**
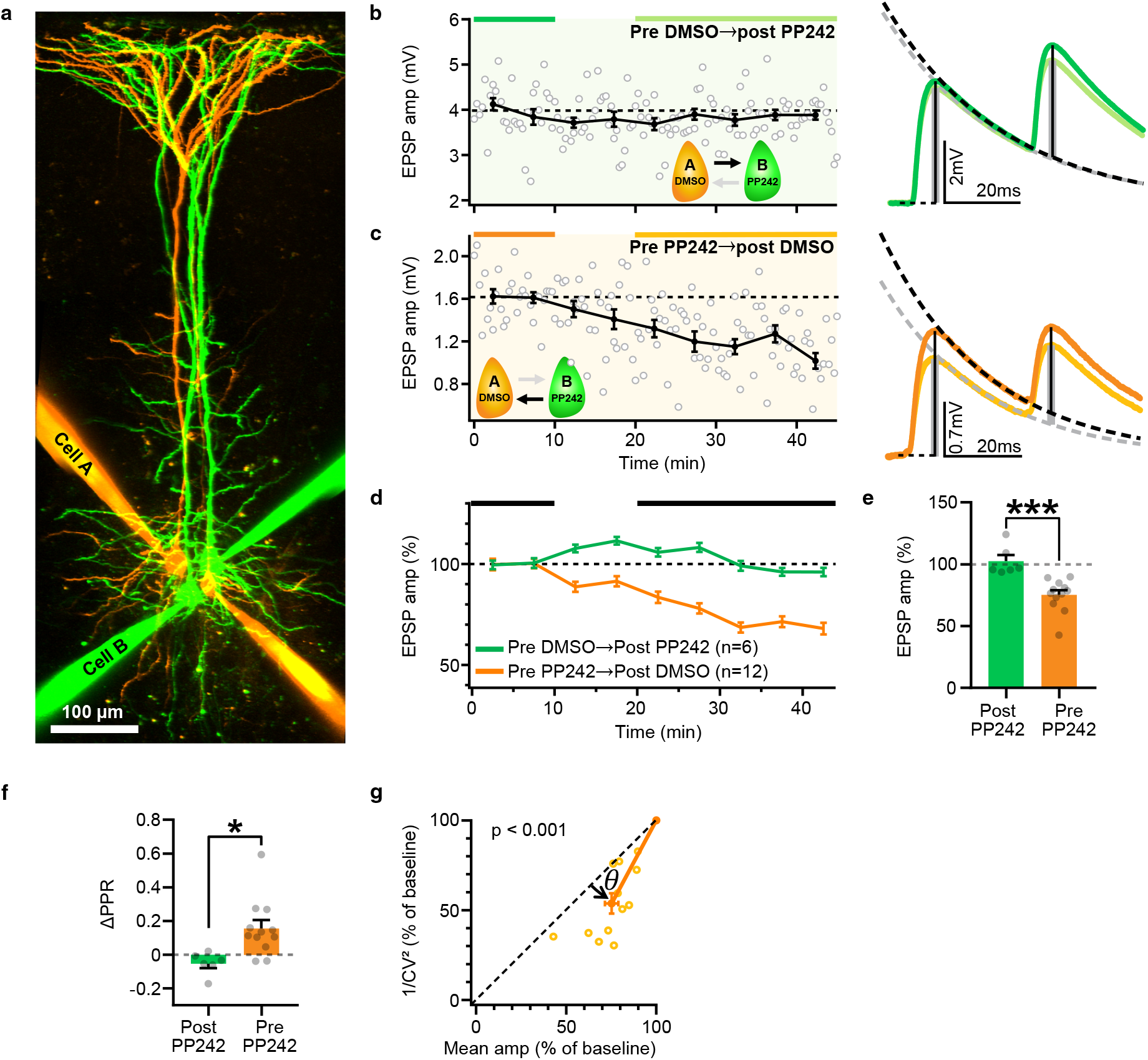
Presynaptic translation supports evoked release via ATP-mediated mTOR signalling. **a,** 2PLSM image of a sample quadruple whole-cell recordings showing bidirectional connections between Cell A (control) and Cell B (PP242-loaded). Scale bar, 100 μm. **b,** Stable EPSPs in connection with postsynaptic PP242 (Cell B). Traces are averaged from and color-coded by the baseline and “after” periods as indicated by bars above the line graph. **c,** Same as (b) for weakened EPSPs in the reciprocal connection with presynaptic PP242 (Cell B). **d,** Ensemble averages. (n = connections). **e,** Presynaptic, but not postsynaptic, PP242 loading reduced EPSP amplitude (*t_16_ = 4.33*, unpaired t test, \*\*\**P* < 0.001). **f,** Presynaptic PP242 loading increased PPR, indicating short-term facilitation (*t_16_ = 2.87*, unpaired t test, \**P* < 0.05). **g,** CV analysis showing points are below the diagonal for connections with presynaptic M7 loading (Wilcoxon test, *θ* = 16° ± 3°, *P* < 0.001), indicating a presynaptic locus of depression. Mean ± s.e.m.

**Fig. S4.**
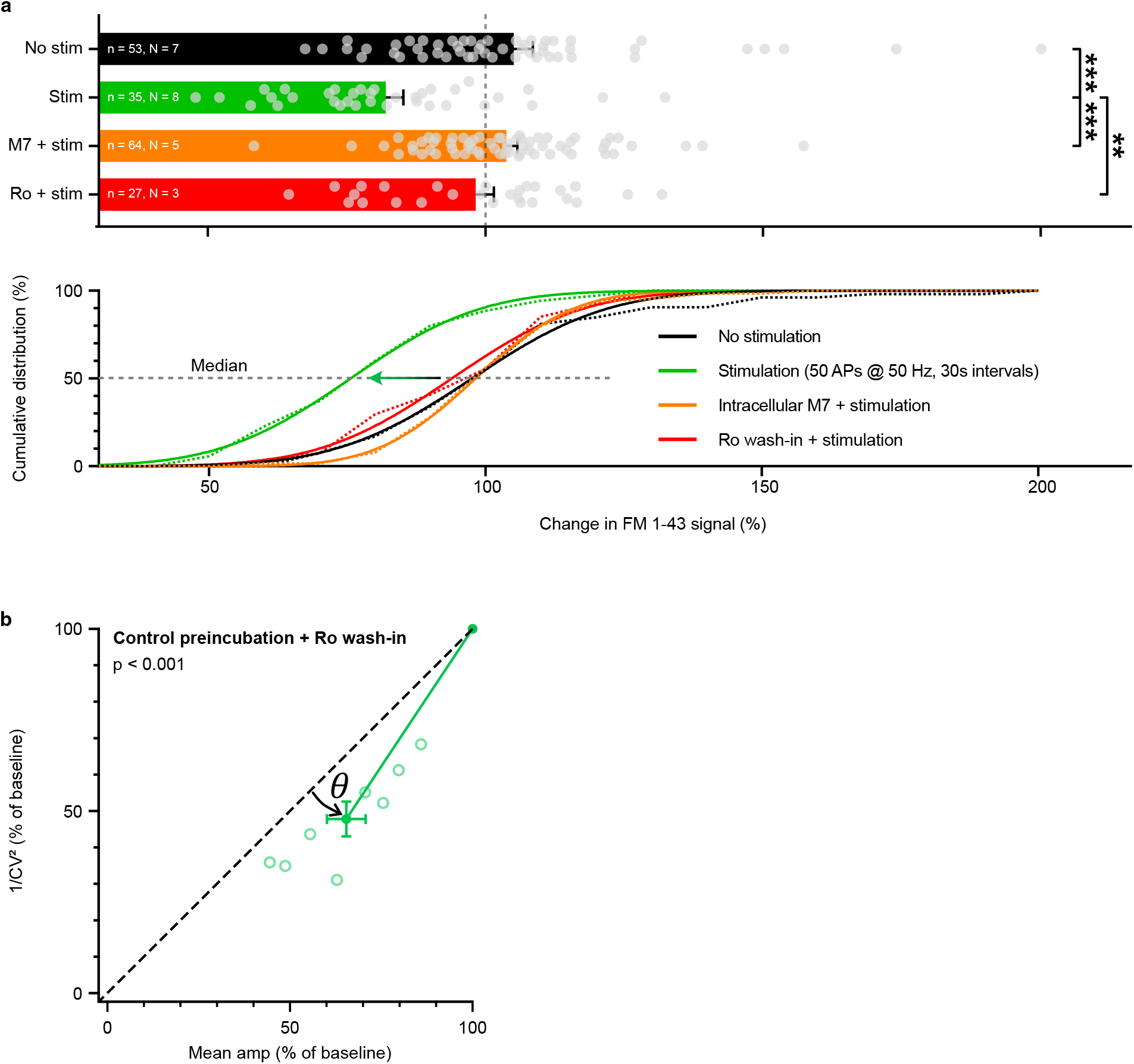
Inhibition of preNMDARs and translation supresses evoked release. **a,** Top: Scatter plot showing the inhibition of preNMDAR (Ro wash-in) or presynaptic translation (intracellular M7 loading) abolished the vesicle release-driven FM 1-43 signal drop (*n = boutons, N = slices;* Kruskal-Wallis test, *P* < 0.001, \*\**P* < 0.01, \*\*\**P* < 0.001 with Dunn’s correction). Bottom: Cumulative distributions showing the data (dotted) and Gaussian fits (solid lines) demonstrate a notable left shift for the stimulation-only condition. **b,** Left: CV analysis showing points are below the diagonal for connections with control preincubation and Ro wash-in (Wilcoxon test, *θ* = 13° ± 2°, *P* < 0.001), indicating a presynaptic locus of expression. CHX preincubation occluded the effect of Ro in CV analysis, consistent with the model that presynaptic translation mediates preNMDAR-dependent synaptic release. Mean ± s.e.m.

**Fig. S5.**
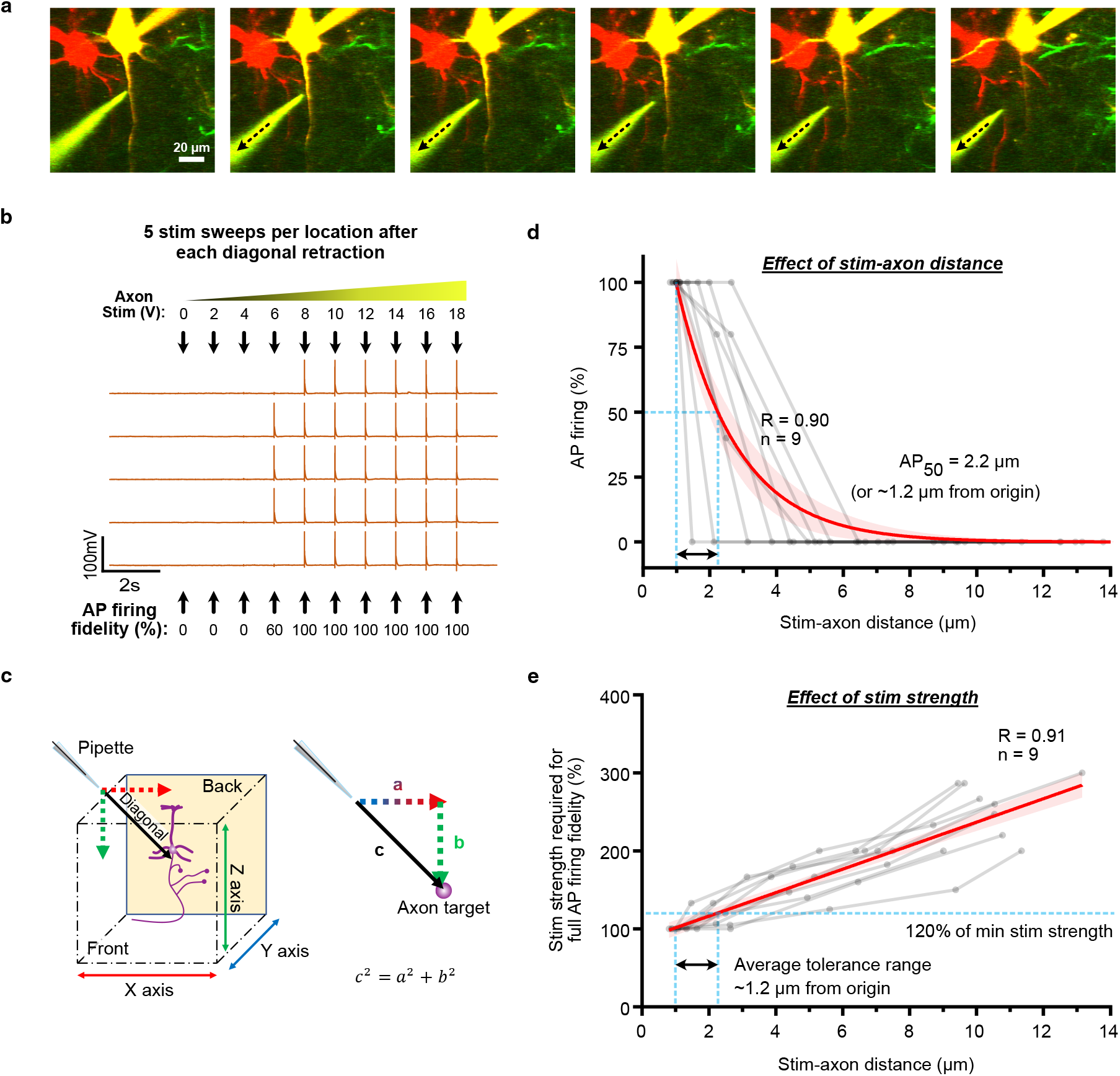
Effect of stim-axon distance and strength on action potential firing. **a,** Single-plane 2PLSM images of a sample experiment with progressively retracted axon stimulation pipette (dotted arrow). The focus was placed on the Z plane with the pipette tip to present a sense of distance from the axon. Scale bar, 20 μm. **b,** At each pipette location as in (**a**), one spike each in a voltage ramp of 0 – 18 V in 2 V steps were applied via the stimulation pipettes with 1s inter-ramp intervals. The sweep was repeated 5 times and the succeed rate of AP firing was extracted for each stimulation voltage. **c,** Pythagoras theorem, *c^2^ = a^2^ + b^2^*, was used to derive the closest diagonal stim-axon distance. **d,** AP firing success rate reduced as stim-axon distancing increased. The data was fitted with exponential decay function (red, with 95% confidence intervals in pink). AP firing success rate was reduced to 50% (AP_50_) when the stimulation pipette was moved ~1.2 μm. **e,** Minimum stimulation strength required for full AP firing fidelity increased as stim-axon distance increased. The data was fitted with line (red, with 95% confidence intervals in pink). On average, 120% of minimum stimulation voltage provides an average spatial tolerance of ~1.2 μm from the stimulation origin.

**Fig. S6.**
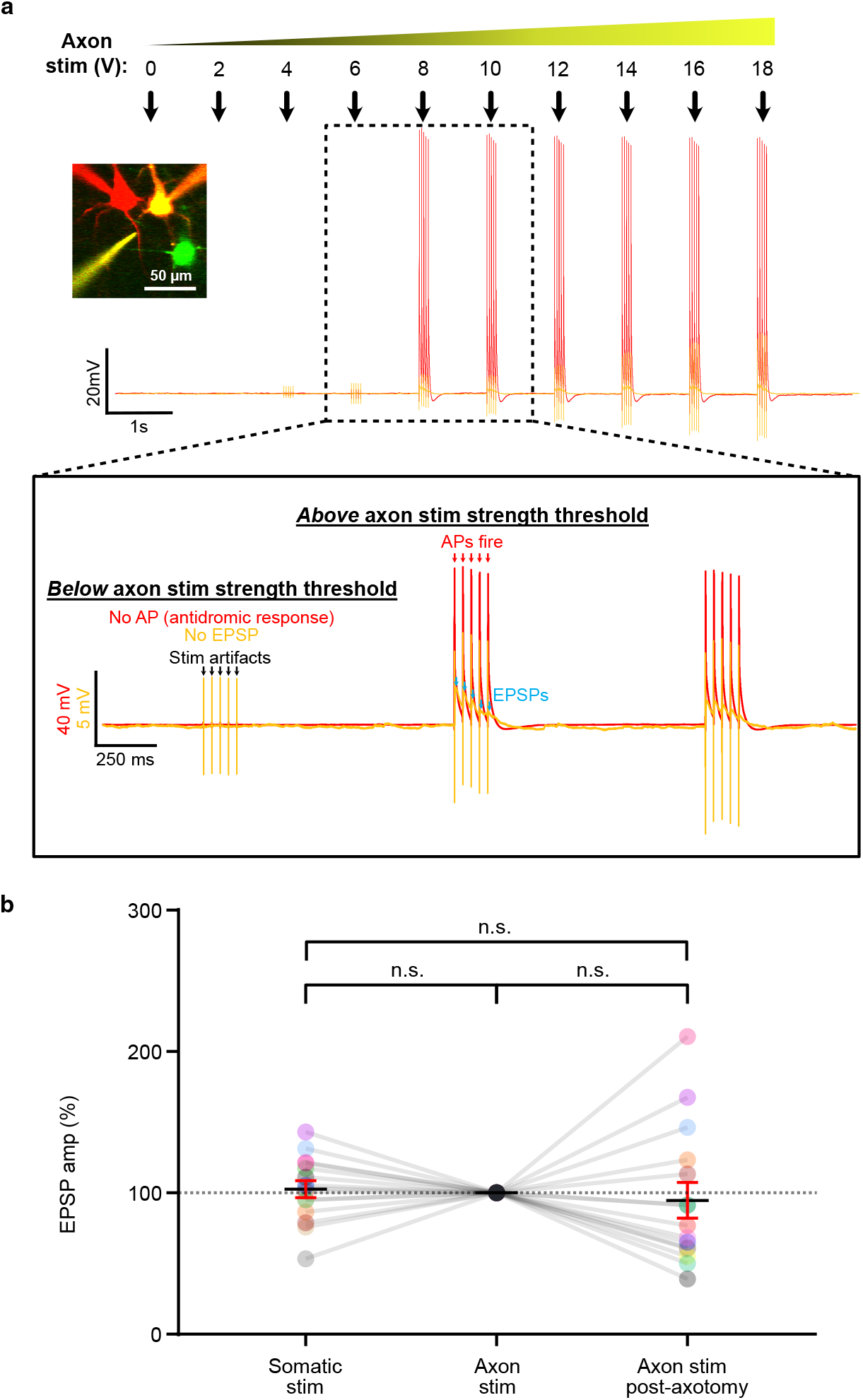
Establishing axon-evoked action potentials and synaptic release. **a,** With 2PLSM and laser Dodt-contrast real-time imaging, the stimulation pipette was guided to the position of ~1 μm from the axon initial segment. Five spikes each in a voltage ramp of 0 – 18 V in 2 V steps were applied via the stimulation pipettes with 1 s inter-ramp intervals. Three to 5 repeats were performed to identify a reliable minimum stimulation voltage threshold required for evoking action potentials (APs; measured as antidromic responses with somatic pipette; red arrows). Millisecond-coupled axon-evoked EPSPs (blue arrows) in the postsynaptic cell should only be observed when the minimum threshold for triggering AP firing is reached. Scale bar, 50 μm. **b,** EPSP amplitudes were indistinguishable with somatic stimulation, axon stimulation pre-axotomy, and axon stimulation post-axotomy (*n = 15 connections, F_1.36, 19.0_ = 0.31*, repeated measures ANOVA, *P* = 0.65). Mean ± s.e.m.

**Fig. S7.**
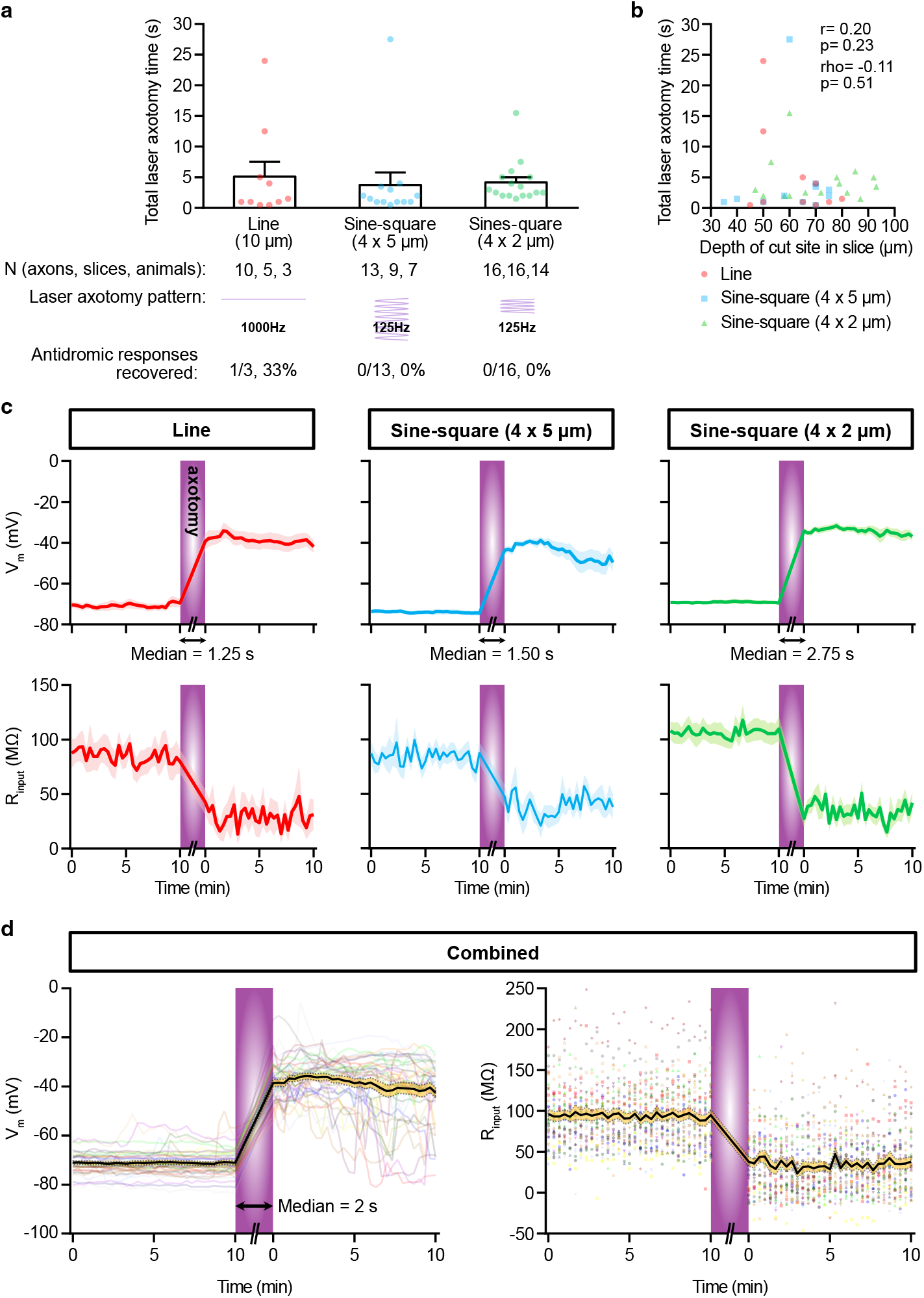
Optimization of 2-photon laser microsurgery for axotomy. **a,** Line, 4 5 μm sine-square, and 4 2 μm sine-square laser cuts are compared. Laser cuts are performed in units of 500 ms every 10s, with laser tuned to 780 nm and an objective back aperture power of 50 mW. Total laser axotomy time were indistinguishable across the three groups. (N = axons/cells, slices, animals; Kruskal-Wallis test, *P* = 0.07). However, 1 out 3 tested neurons (33%) in the line cut condition showed recovery of antidromic responses, suggesting partial recovery. **b,** Laser microsurgery was efficient down to ~100 μm, the deepest tested (r = 0.20, *P* = 0.23; rho = 0.11, *P* = 0.51). **c,** Successful laser axotomy was accompanied by rapid depolarization of membrane potential (V_m_) and input resistance (R_input_). **d,** V_m_ and R_input_ changes are plotted as individual (multi-colour) and ensemble average traces (black lines with s.e.m. in orange shade) pooled from all three cutting methods. Median axotomy time was 2 s (n=39). In 12 out of the 39 samples, traces of voltage ramps without the test pulses needed for R_input_ measurement are omitted in the R_input_ graph. Mean ± s.e.m.

**Fig. S8.**
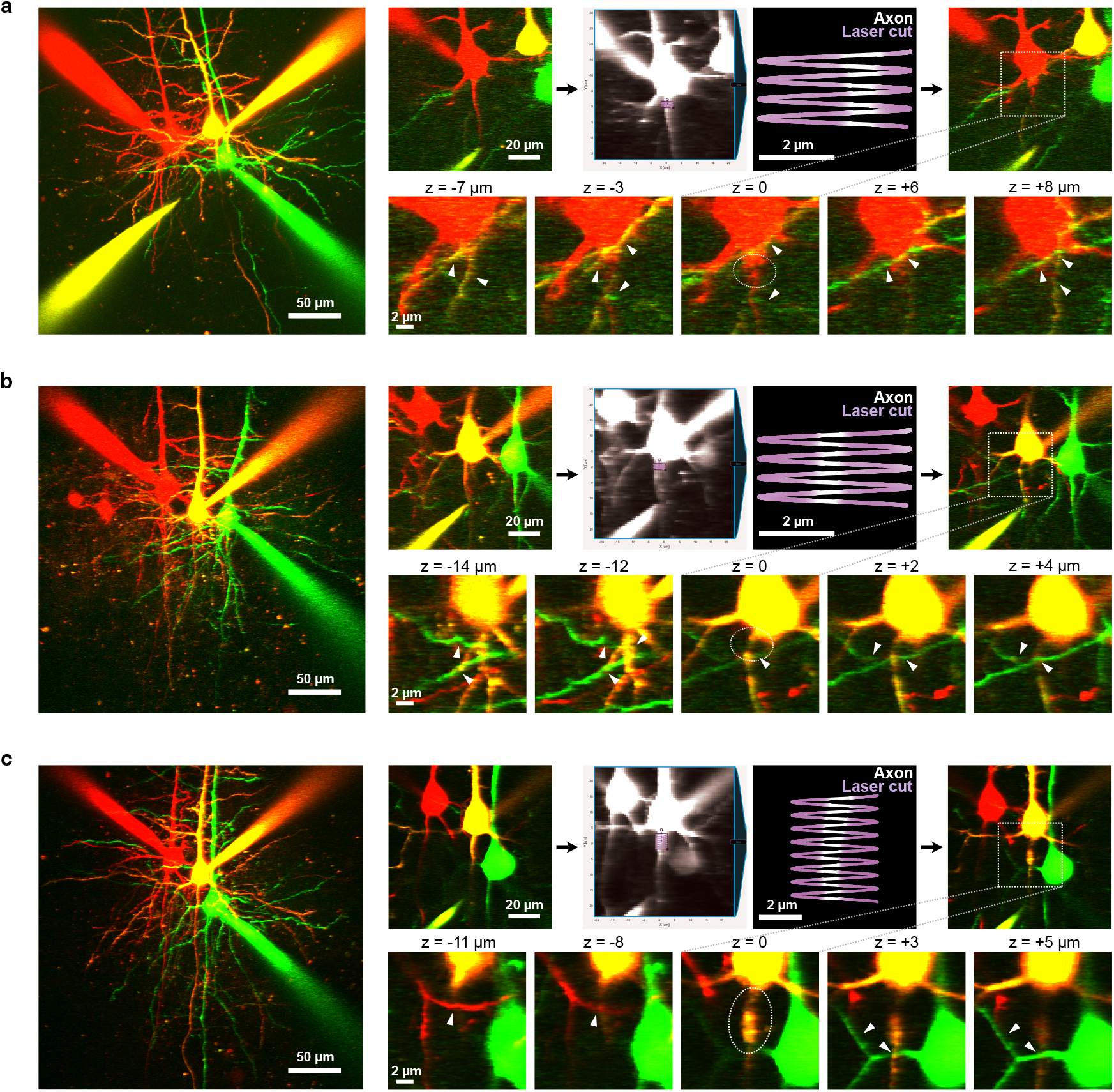
Two-photon laser microsurgery offers micron precision. **a,** Sample triplet-recordings with cell in red targeted for 4 2 μm sines-quare axotomy. Dotted area is enlarged in the second row. Note the neurites within microns in the x, y and z planes (white arrow heads) were intact after axotomy. **b,** Same as (**a**) with cell in orange targeted for 4 2 μm sinesquare axotomy. **c,** Same as (**a**) with cell in orange targeted for 4 5 μm sine-square axotomy. Scale bars, 50 μm for images on left, 20 μm for 3 zoom, 2 μm for axotomy image and images in the second row.

**Fig. S9.**
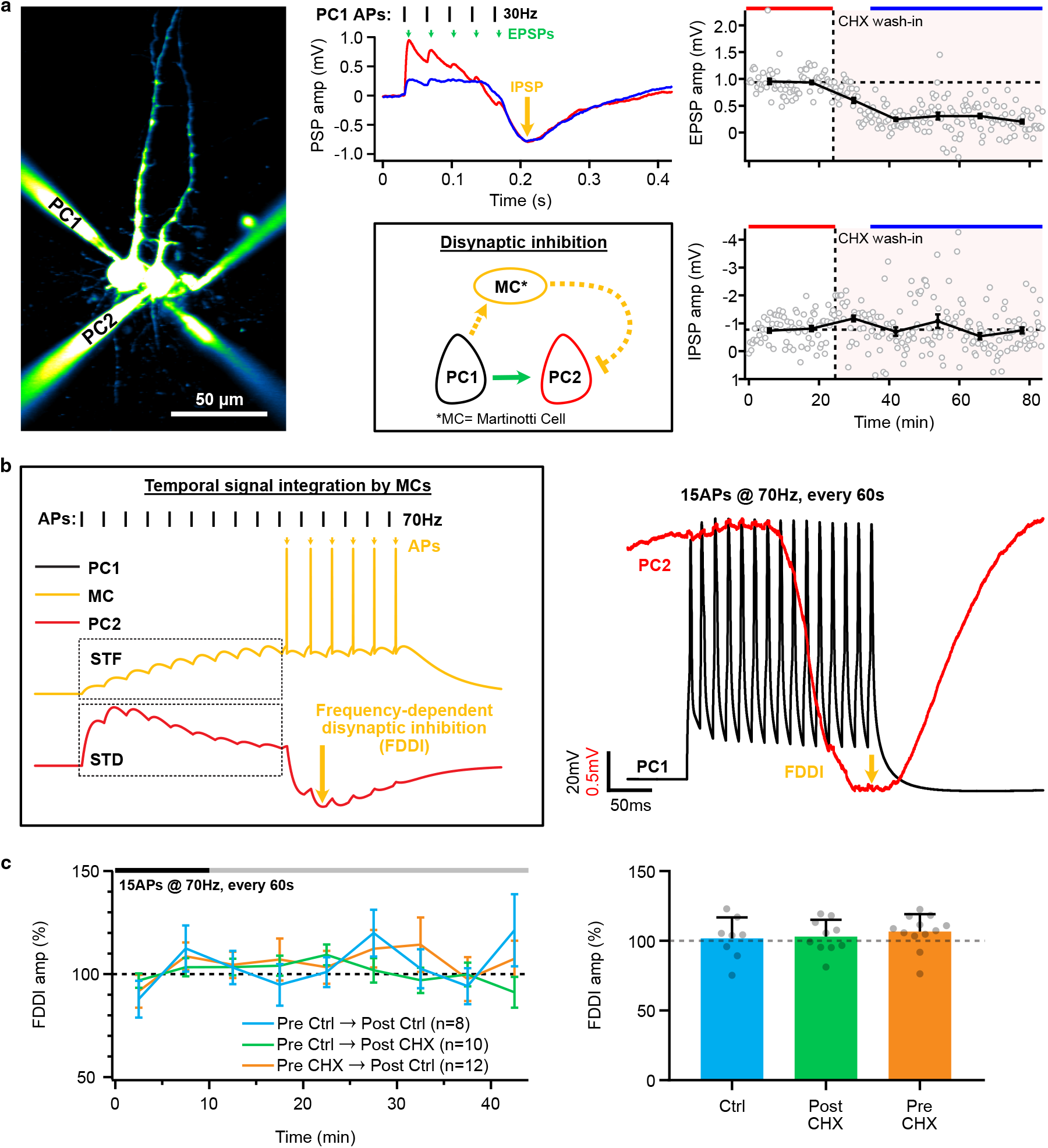
Presynaptic translation inhibition does not affect frequency-dependent disynaptic inhibition. **a,** Sample whole-cell recordings showing two types of connections between PC1 and PC2 at 30 Hz stimulation: monosynaptic EPSPs and disynaptic IPSP. While CHX wash-in acutely inhibited EPSP amplitude (green arrows), the delayed inhibitory component (IPSPs; orange arrows) was unaffected. Traces are averaged from and color-coded by the baseline and “after” periods as indicated by bars above the line graph. Scale bar, 50 μm. **b,** Simulation showing the monosynaptic PC1→PC2 excitatory connection and the disynaptic PC1→MC→PC2 circuit motif in (**a**). PC→MC connections are strongly short-term facilitating, in contrast to PC→PC connections, which are short-term depressing. MC only fires once the temporally summating EPSPs reach spike threshold, thus resulting in a delayed inhibitory component in the postsynaptic PC. This phenomenon occurs reliably when presynaptic PC fires at high frequency, such as with the 15 spikes at 70 Hz stimulation protocol^33,41,42^. This is therefore known as frequency-dependent disynaptic inhibition (FDDI; orange arrows). **c,** Ensemble averages show that FDDI amplitudes were unaffected by pre- or postsynaptic CHX loading (n = connections, *F_2, 27_ = 0.38*, one-way ANOVA, *P* = 0.69), in contrast to the suppression of EPSP amplitude by presynaptic CHX (Fig. S1). Mean ± s.e.m.

**Fig. S10.**
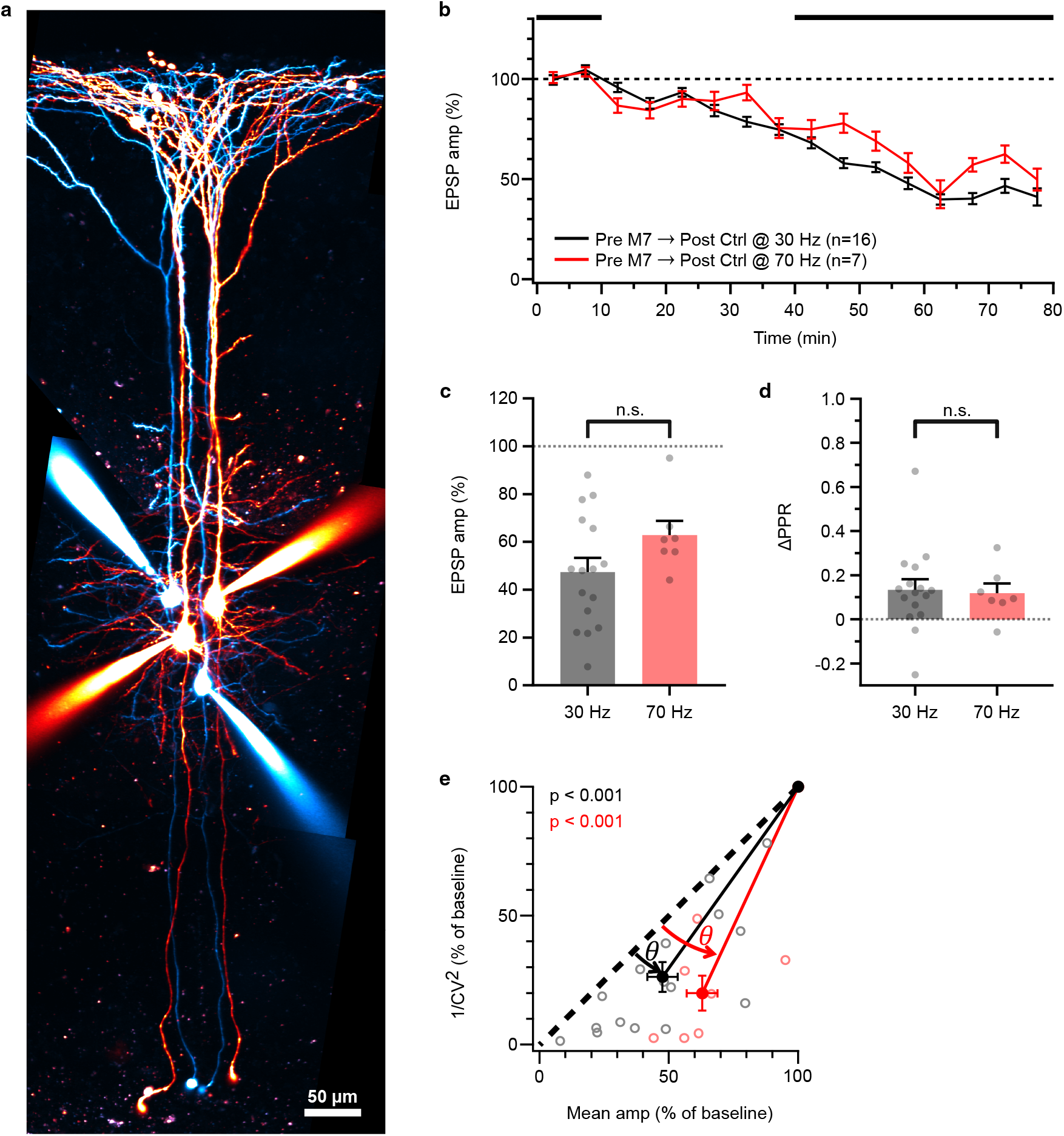
Presynaptic translation sustains PC→PC synaptic release at both 30 Hz and 70 Hz. **a,** 2PLSM image of layer-5 PCs in intracellular M7 loading experiment. Scale bar, 50 μm. **b,** Ensemble averages showing similar suppression of PC→PC synapses at both 30 Hz and 70 Hz. (n = connections). 30-Hz frequency data were reanalysed (Fig. 1d-j). **c,** PC→PC EPSP amplitudes were indistinguishable (*t_21_ = 1.58*, unpaired t test, *P* = 0.13). **d,** PPRs were indistinguishable (*t_21_ = 0.19*, unpaired t test, *P* = 0.85). **e,** CV analysis showed points below the diagonal for both stimulation frequencies (Wilcoxon test, 30 Hz: *θ* = 10° ± 2°, *P* < 0.001; 70 Hz: *θ* = 20° ±4°, *P* < 0.001), indicating presynaptic loci of depression. Mean ± s.e.m.

